# Cancer cell glycocalyx thickness is a druggable physical barrier to cellular immunotherapy

**DOI:** 10.1101/2024.12.05.627089

**Authors:** Sangwoo Park, Justin H. Paek, Marshall J. Colville, Ling-Ting Huang, Audrey P. Struzyk, Sriram Neelamegham, Marcela V. Maus, Matthew J. Paszek

## Abstract

The abnormally thick glycocalyx of cancer cells can physically limit immune-cell engagement and reduce the efficacy of cellular immunotherapies, but strategies for identifying pharmacological agents that relieve this barrier remain limited. Here, we establish glycocalyx thickness as a pharmacologically actionable physical phenotype. Disruption of mucin-type *O*-glycosylation reduced glycocalyx thickness and increased NK- and CAR-NK-mediated killing, demonstrating that pharmacological glycocalyx remodeling can enhance innate and CAR-directed cytotoxicity. To screen this phenotype directly, we developed a scanning angle interference microscopy (SAIM) platform using complementary leucine zipper pairs as a stable extracellular plasma-membrane reference. A focused screen of glycosylation and metabolic inhibitors resolved compound- and cell-type-specific nanoscale changes in glycocalyx thickness and identified several thinning agents. Castanospermine, an inhibitor of early *N*-glycan processing, was among the strongest hits and reduced mucin-probe binding in an independent mucin-rich gastric cancer model. Castanospermine-treated gastric cancer cells retained MUC17 antibody recognition and showed increased physical avidity for MUC17 CAR-T cells, while castanospermine enhanced MUC17 CAR-T-mediated tumor control in real-time co-culture assays. These findings establish direct glycocalyx-thickness measurement as a physical-phenotype-guided screening strategy and identify castanospermine as a candidate modulator for combination with MUC17 CAR-T cells.

## Introduction

The cellular glycocalyx is a dense meshwork of glycosylated macromolecules that covers the surface of all eukaryotic cells. As the first point of contact between interacting cells, the glycocalyx constitutes an important component of cell-cell interfaces, including immune synapses formed between cytotoxic immune cells and target cancer cells^1,2^. Malignant transformation is commonly accompanied by dysregulated glycosylation and remodeling of the cell-surface glycan landscape^3–5^. In addition, many tumors overexpress bulky and highly hydrated glycopolymers, including cell-surface mucins and glycosaminoglycans, which can markedly increase the physical thickness of the glycocalyx^1,2^. Therefore cancer cells can exhibit major changes in both the molecular composition and physical structure of the glycocalyx, creating a protective cell-surface layer that contributes to immune evasion^2,6–8^.

Efforts to therapeutically target the cancer glycocalyx have largely focused on its biochemical functions. Tumor-associated glycans can engage carbohydrate-binding receptors on immune cells and suppress antitumor immune responses, leading to the development of strategies that inhibit the biosynthesis of glyco-immune checkpoint ligands, block their cognate receptors, or enzymatically remove cancer-associated glycoproteins and mucins^7–10^. However, biochemical signaling represents only one mechanism by which the glycocalyx can regulate immune-cell function. Our previous work demonstrated that a dense glycocalyx generated by overexpression of the cell-surface mucin MUC1 can act as a physical barrier to immune-cell attack, protecting cancer cells even when immune cells are engineered with chimeric antigen receptors (CARs) that provide strong antigen-specific activation^2^. Importantly, nanometer-scale changes in glycocalyx thickness were sufficient to substantially alter susceptibility to immune-cell-mediated killing. These findings suggest that the physical dimensions of the cancer glycocalyx represent an additional therapeutic parameter that could be targeted to improve cellular immunotherapy.

One approach to overcome this physical barrier is to reduce the abundance or structural extension of glycocalyx components. Mucin-selective proteases such as StcE can remove bulky cell surface mucins^11,12^. Similarly, pharmacological inhibition of tumor N-glycosylation with 2-deoxy-D-glucose has been shown to enhance CAR-T-cell activity across multiple solid tumor models^13^. However, pharmacological perturbations are generally selected on the basis of their annotated biochemical targets^14–16^. The effects of these perturbations on the physical thickness of the glycocalyx are difficult to predict. Because the glycocalyx composition differs across cancer cell types, inhibition of the same glycosylation pathway may have different effects on glycocalyx structure^1,17^. A method that directly measures the physical structure of the glycocalyx could provide a useful phenotypic readout for identifying compounds that physically remodel the cancer-cell surface, rather than relying solely on the annotated biochemical target of an inhibitor.

The precise measurement of the physical thickness of the glycocalyx in live cells at the nanometer scale remains difficult. Super-resolution approaches have demonstrated that changes associated with malignant transformation, including oncogenic signaling and epithelial-to-mesenchymal transition, can substantially alter glycocalyx height^18^. However, many high-resolution imaging methods require long acquisition times or sample fixation, limiting their application to live-cell pharmacological screening^19^. To address this need, our group developed scanning angle interference microscopy (SAIM), an optical method that enables nanoscale axial localization of fluorescent probes and quantitative measurement of glycocalyx thickness^2,20–23^. SAIM measures the axial position of fluorophores through interference generated between incident and reflected excitation light, providing rapid nanoscale localization compatible with live-cell imaging. However, for glycocalyx measurements, SAIM requires a bright, specific, and stable fluorescent reference at the plasma membrane. Conventional strategies present several limitations. Genetically encoded fluorescent membrane proteins can accumulate within intracellular trafficking compartments^24^, whereas lipophilic membrane dyes can redistribute from the plasma membrane, undergo endocytosis, and exhibit cell type-dependent labeling efficiency^25,26^. These limitations become particularly important when SAIM is applied across multiple cancer cell lines or over extended imaging periods.

Here, we use cancer-cell glycocalyx thickness as a physical phenotype for pharmacological screening. We developed a complementary leucine-zipper membrane reference and integrated it with SAIM to measure nanoscale changes in glycocalyx thickness across different cancer cell types^27–29^. Using this platform, we screened a focused panel of pharmacological compounds and identified several agents that reduced glycocalyx thickness, with substantial differences across cell types. We then combined the SAIM screen with mucin and lectin profiling in GSU gastric cancer cells, which led us to prioritize castanospermine for further testing. Pretreatment of GSU cells with castanospermine increased their avidity for MUC17 CAR-T cells, and combining castanospermine with MUC17 CAR-T cells improved tumor control in co-culture. Together, these results show that direct measurement of glycocalyx thickness can be used to identify pharmacological compounds with potential to improve cellular immunotherapy.

## Results

### Pharmacological reduction of glycocalyx thickness enhances immune-cell cytotoxicity

We previously demonstrated that the cancer-cell glycocalyx can function as a physical barrier that limits productive interactions between cancer cells and immune cells, thereby reducing immune-cell-mediated cytotoxicity^2^. Bulky cell-surface glycoproteins, including mucins such as MUC1 as well as other extended glycoproteins such as CD43, can contribute to this steric barrier and restrict close cell-cell contact^30,31^. We and others have also shown that enzymatic removal of cell-surface glycoproteins or glycans, including mucin degradation by StcE, can reduce glycocalyx thickness and improve immune-cell access to the target-cell surface^6,9–11,32^. Based on these observations, we hypothesized that pharmacological compounds capable of reducing glycocalyx thickness could similarly enhance immune cell-mediated killing (**Fig. 1a,b**). To test this hypothesis, we first used Ac_5_GalNTGc, a metabolic inhibitor of mucin-type *O*-glycosylation, in a previously established MCF10A-derived cell line with doxycycline-inducible MUC1 expression (1E7)^2,14,33,34^. In this system, mOxGFP is inserted between the MUC1 ectodomain and transmembrane domain, allowing cell-surface MUC1 abundance to be directly visualized across a tunable range of MUC1 expression. Increasing doxycycline concentrations progressively increased MUC1 expression, whereas treatment with Ac_5_GalNTGc reduced the MUC1-associated fluorescence signal, particularly at higher levels of MUC1 induction (**Fig. 1c**). Consistent with inhibition of core-1 *O*-glycan elaboration, Ac_5_GalNTGc treatment also reduced PNA lectin binding across the doxycycline concentrations tested^14^ (**Fig. 1d**).

**Figure 1.**
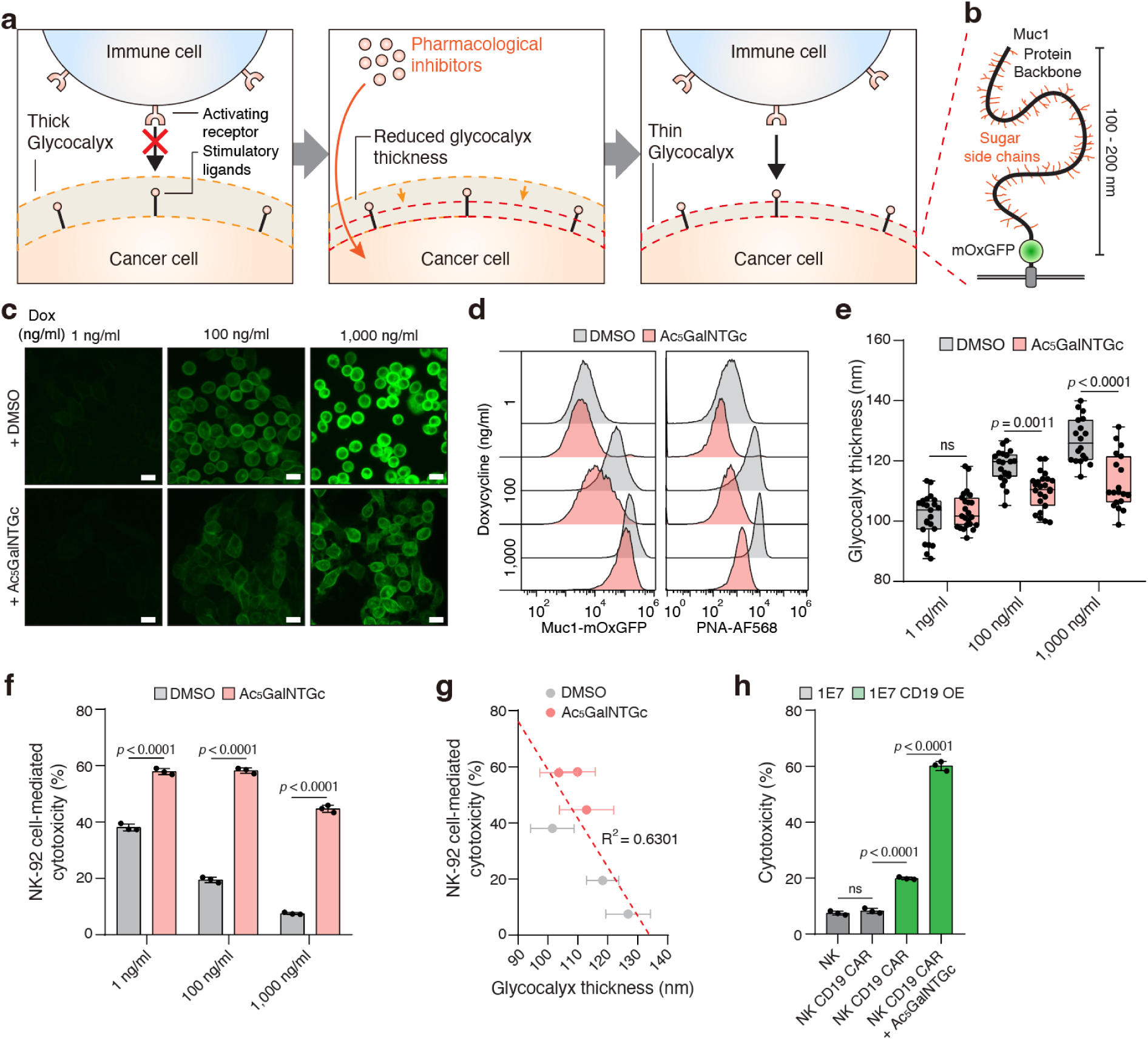
Glycocalyx thickness is a functional barrier to immune-cell cytotoxicity and can be pharmacologically modulated. **a**, Schematic illustrating how a thick cancer-cell glycocalyx can limit productive immune-cell engagement and how pharmacological inhibitors or targeted degradation strategies may reduce glycocalyx thickness to improve receptor accessibility and immune-cell recognition. **b**, Schematic representation of cell-surface MUC1, a major glycocalyx component consisting of a membrane-anchored protein backbone decorated with extended glycan side chains. **c**, Representative fluorescence images of the doxycycline-inducible MUC1 1E7 clone cultured with the indicated concentrations of doxycycline (1, 100, or 1,000 ng/ml) and treated with DMSO or 60 μM Ac_5_GalNTGc. Scale bar, 200 µm. **d**, Flow cytometry analysis of cell-surface MUC1 (MUC1-mOxGFP) and PNA-AF568 binding in 1E7 cells cultured with the indicated doxycycline concentrations following treatment with DMSO or 100 μM Ac_5_GalNTGc. **e**, Quantification of glycocalyx thickness in 1E7 cells cultured with the indicated doxycycline concentrations following treatment with DMSO or Ac_5_GalNTGc. Boxes indicate the first and third quartiles, center lines indicate the median, and whiskers indicate the range. **f**, NK-92 cell-mediated cytotoxicity against 1E7 target cells expressing the indicated levels of MUC1 after treatment with DMSO or Ac_5_GalNTGc. Cytotoxicity was measured after 4h of co-culture. NK-cell-to-target-cell ratio is 5:1. Results are mean ± s.d. of n = 3. **g**, Correlation between NK-92 cell cytotoxicity and glycocalyx thickness (R² = 0.6301); data from panels **e** and **f**; The dashed line represents a linear regression. **h**, Cytotoxicity of unmodified NK-92 cells or CD19 CAR-expressing NK-92 cells against parental 1E7 cells or 1E7 cells overexpressing CD19, with or without Ac_5_GalNTGc treatment. In **e,f,h**, statistical analysis was performed by one-way ANOVA with Tukey’s post hoc tests.

We next asked whether these changes in cell-surface mucins and glycans translated into measurable changes in the physical structure of the glycocalyx. Using Scanning Angle Interference Microscopy (SAIM) to quantify glycocalyx thickness, we found that the effect of Ac_5_GalNTGc depended strongly on MUC1 abundance. At low doxycycline concentrations, where MUC1 expression was limited, glycocalyx thickness was similar between DMSO- and Ac_5_GalNTGc-treated cells. In contrast, the difference became progressively greater as MUC1 expression increased, with Ac_5_GalNTGc reducing glycocalyx thickness by 13.9 nm relative to DMSO-treated cells at maximal MUC1 induction (**Fig. 1e**). An independent experiment reproduced this MUC1-dependent reduction in glycocalyx thickness (**Supplementary Fig. 1a**). These results indicate that pharmacological disruption of mucin glycosylation can substantially reduce the physical dimensions of a mucin-rich glycocalyx.

We then determined whether this reduction in glycocalyx thickness was sufficient to alter susceptibility to immune-cell attack. 1E7 target cells were treated with Ac_5_GalNTGc for 48 h and subsequently co-cultured with NK-92 cells for 4 h. Ac_5_GalNTGc treatment substantially increased NK-92-mediated cytotoxicity, with the largest enhancement observed under conditions of high MUC1 expression (**Fig. 1f**). Across the different levels of MUC1 induction, glycocalyx thickness was inversely associated with NK-92-mediated cytotoxicity (**Fig. 1g**), consistent with a model in which a thicker glycocalyx provides greater protection from immune-cell attack^2^. Finally, to test whether pharmacological glycocalyx reduction could also enhance the activity of engineered immune cells, we generated 1E7 cells overexpressing CD19 and compared the cytotoxic activity of unmodified NK-92 cells and previously established CD19 CAR-NK-92 cells in the presence or absence of Ac_5_GalNTGc^2^. CD19 expression alone increased susceptibility to CD19 CAR-NK-mediated killing, whereas combining CD19 CAR-NK cells with Ac_5_GalNTGc treatment produced a 3.02-fold increase in cytotoxicity compared with CD19 CAR-NK cells against untreated target cells (**Fig. 1h**). Together, these results establish that pharmacological reduction of the cancer cell glycocalyx can enhance both innate and CAR-directed immune-cell cytotoxicity and support the use of glycocalyx thickness as a pharmacologically actionable phenotype for improving cellular immunotherapy.

### A leucine zipper-based strategy enables stable plasma-membrane labeling for SAIM

The ability of pharmacological glycocalyx modulation to enhance immune-cell cytotoxicity prompted us to develop a scalable approach for directly measuring glycocalyx thickness as a phenotypic readout for drug screening. SAIM enables nanoscale axial localization of fluorescent probes and can therefore be used to quantify the physical dimensions of the cell-surface glycocalyx. Accurate measurement of glycocalyx thickness without artifacts introduced by chemical fixation requires a bright and stable fluorescent reference to the plasma membrane in live cells. In our initial experiments (**Fig. 1e**), we used both the extracellular mOxGFP tag in the engineered 1E7 MUC1 model and an optimized commercial plasma-membrane dye as reference markers for SAIM, and obtained comparable measurements with the two approaches. The mOxGFP tag provided a bright membrane reference, but fluorescence from intracellular protein in the ER and Golgi could interfere with fitting of the plasma-membrane signal. Commercial membrane dyes avoided this intracellular fluorescent pool, but required substantial optimization for each cell line. In some conditions, prolonged live-cell imaging also resulted in loss of membrane-associated fluorescence, intracellular accumulation, reduced cell viability, or cell detachment^35,36^. Due to these limitations, we sought a membrane-labeling strategy that would provide bright and stable fluorescence specifically at the extracellular plasma membrane.

To achieve this, we developed a labeling system based on a pair of engineered leucine zipper peptides, referred to here as AZip and BZip, which associate with low-nanomolar affinity through complementary electrostatic and hydrophobic interactions^27^. BZip was genetically anchored to the plasma membrane through the PDGFRβ transmembrane domain^37^, whereas recombinant AZip served as an extracellular fluorescent labeling probe (**Fig. 2a,b**). We initially generated AZip fused to superfolder GFP (sfGFP) and tested labeling in BZip-expressing cancer cells. AZip-sfGFP produced bright plasma-membrane labeling in Capan-2 pancreatic cancer cells (**Fig. 2c**) and across multiple breast cancer cell lines, including KPL-1, T47D, SKBR3, and ZR-75-1 (**Fig. 2d**). In contrast, direct fusion of mScarlet to membrane-anchored BZip resulted in substantial intracellular fluorescence, whereas extracellular labeling with AZip-sfGFP more selectively delineated the cell surface (**Fig. 2d**).

**Figure 2.**
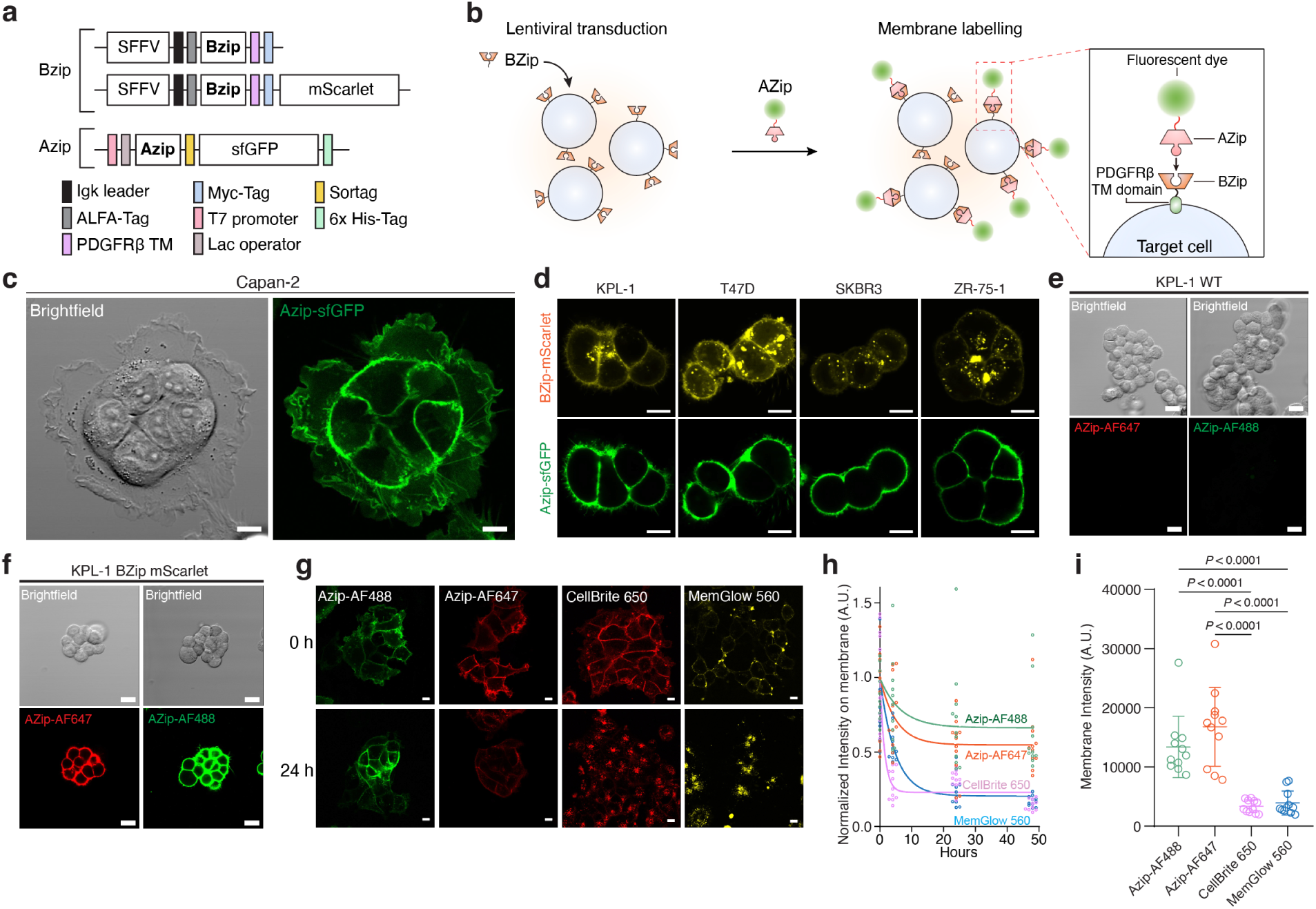
Validation of the leucine zipper-based membrane labelling strategy. **a**, Schematic representation of constructs used for expression of membrane-anchored BZip, BZip-mScarlet, and AZip-sfGFP. BZip constructs contain a PDGFRβ transmembrane (TM) domain for cell-surface anchoring. **b**, Schematic of cell-surface labeling through the AZip-BZip interaction. Target cells are transduced to express membrane-anchored BZip and subsequently labeled with fluorescently tagged AZip. **c**, Representative bright-field and fluorescence images of BZip-expressing Capan-2 pancreatic cancer cells labeled with 1 μM AZip-sfGFP for 1 h at 4 °C. Scale bars, 10 μm. **d**, Representative fluorescence images of BZip-mScarlet expression and AZip-sfGFP labeling in the indicated cancer cell lines after incubation with AZip-sfGFP for 1 h at 4 °C. Scale bars, 10 μm. **e,f**, Representative bright-field and fluorescence images of wild-type KPL-1 cells (**e**) and BZip-mScarlet-expressing KPL-1 cells (**f**) following labeling with AZip-AF488 or AZip-AF647. Scale bars, 10 μm. **g**, Representative fluorescence images of BZip-expressing KPL-1 cells immediately after labeling (0 h) and 24 h after labeling with AZip-AF488, AZip-AF647, CellBrite 650, or MemGlow 560. Cells were labeled according to the indicated conditions and washed twice with culture medium before imaging. Scale bars, 10 μm. **h**, Quantification of normalized membrane fluorescence intensity over time for the indicated membrane-labeling reagents. Membrane fluorescence was quantified using 20 x 20-pixel regions (2.98 x 2.98 μm) from a minimum of 11 cells per condition. **i**, Membrane fluorescence intensity measured immediately after labeling (0 h) with the indicated reagents. In **i**, Statistical significance was determined by one-way ANOVA followed by Tukey’s multiple-comparisons test.

We next adapted the AZip probe for conjugation to small organic fluorophores, including Alexa Fluor 488 (AF488) and Alexa Fluor 647 (AF647), using sortase-mediated ligation (**Supplementary Fig. 2a,b**). This approach reduced the size of the fluorescent labeling probe relative to the sfGFP fusion and provided a compact membrane label suitable for SAIM measurements. Both AZip-AF488 and AZip-AF647 showed minimal labeling of wild-type KPL-1 cells but robust membrane fluorescence in BZip-expressing KPL-1 cells, demonstrating that labeling was dependent on the AZip-BZip interaction (**Fig. 2e,f**). Because SAIM measurements require membrane fluorescence to remain spatially restricted during image acquisition^22,23^, we next compared the stability of AZip-based labeling with commonly used membrane dyes. BZip-expressing KPL-1 cells were labeled with AZip-AF488, AZip-AF647, CellBrite 650, or MemGlow 560 and monitored over time (**Fig. 2g and Supplementary Fig. 2c**). AZip-AF488 and AZip-AF647 exhibited substantially greater retention of fluorescence at the plasma membrane than the conventional membrane dyes, which rapidly lost membrane-associated signal and accumulated intracellularly (**Fig. 2g,h**). AZip-based probes also produced significantly greater membrane fluorescence immediately after labeling than CellBrite 650 or MemGlow 560 (**Fig. 2i**). Among the probes tested, AZip-AF488 showed particularly stable membrane localization over the imaging period. Together, these results establish the AZip-BZip system as a specific and stable membrane-labeling strategy that provides a robust fluorescent reference for subsequent SAIM-based measurements of cancer-cell glycocalyx thickness.

### Leucine zipper-based SAIM detects pharmacologically induced nanoscale remodeling of the cancer-cell glycocalyx

Having established a stable leucine zipper-based membrane-labeling strategy, we next asked whether this approach could be integrated with SAIM to directly quantify nanoscale changes in the cancer-cell glycocalyx. We therefore incorporated AZip-BZip membrane labeling into SAIM as a stable extracellular membrane reference for glycocalyx measurements (**Fig. 3a-c**). Although the inducible MUC1 system provided a controlled model for establishing the relationship between mucin expression and the physical structure of the glycocalyx, we sought to evaluate the platform in a cancer cell line with an endogenous, mucin-rich glycocalyx and without reliance on an engineered MUC1 reporter. Therefore, we checked MUC1 expression across cancer cell lines using publicly available DepMap data to identify a suitable model for pharmacological perturbation studies. Cholangiocarcinoma, pancreatic cancer, and gastric cancer cell lines were among the cancer types with relatively high MUC1 transcript abundance (**Supplementary Fig. 3a**). We next compared MUC1 transcript abundance with cell-surface MUC1 levels measured by flow cytometry across a panel of cancer cell lines. Surface MUC1 measurements were normalized to SKBR3 cells included as an internal reference across experiments. MUC1 transcript abundance was strongly associated with normalized cell-surface MUC1 levels (R² = 0.7768), with the intrahepatic cholangiocarcinoma cell line YSCCC exhibiting particularly high MUC1 surface abundance (**Fig. 3d**). We therefore selected YSCCC as a primary model for evaluating pharmacological changes in the physical structure of the glycocalyx. To establish that the abundant surface mucin layer in YSCCC cells was experimentally modifiable, we treated cells with the mucin-selective protease StcE. StcE treatment substantially reduced cell-surface MUC1 staining (**Fig. 3e**), consistent with enzymatic removal of the mucin-rich glycocalyx. In parallel, reduction of the glycocalyx altered cell-cell and cell-substrate interactions, promoting multicellular aggregate compaction and cell spreading in orthogonal assays (**Extended Data Figs. 4 and 5**). These findings further supported YSCCC as a suitable model in which to quantify physical remodeling of a bulky glycocalyx.

**Figure 3.**
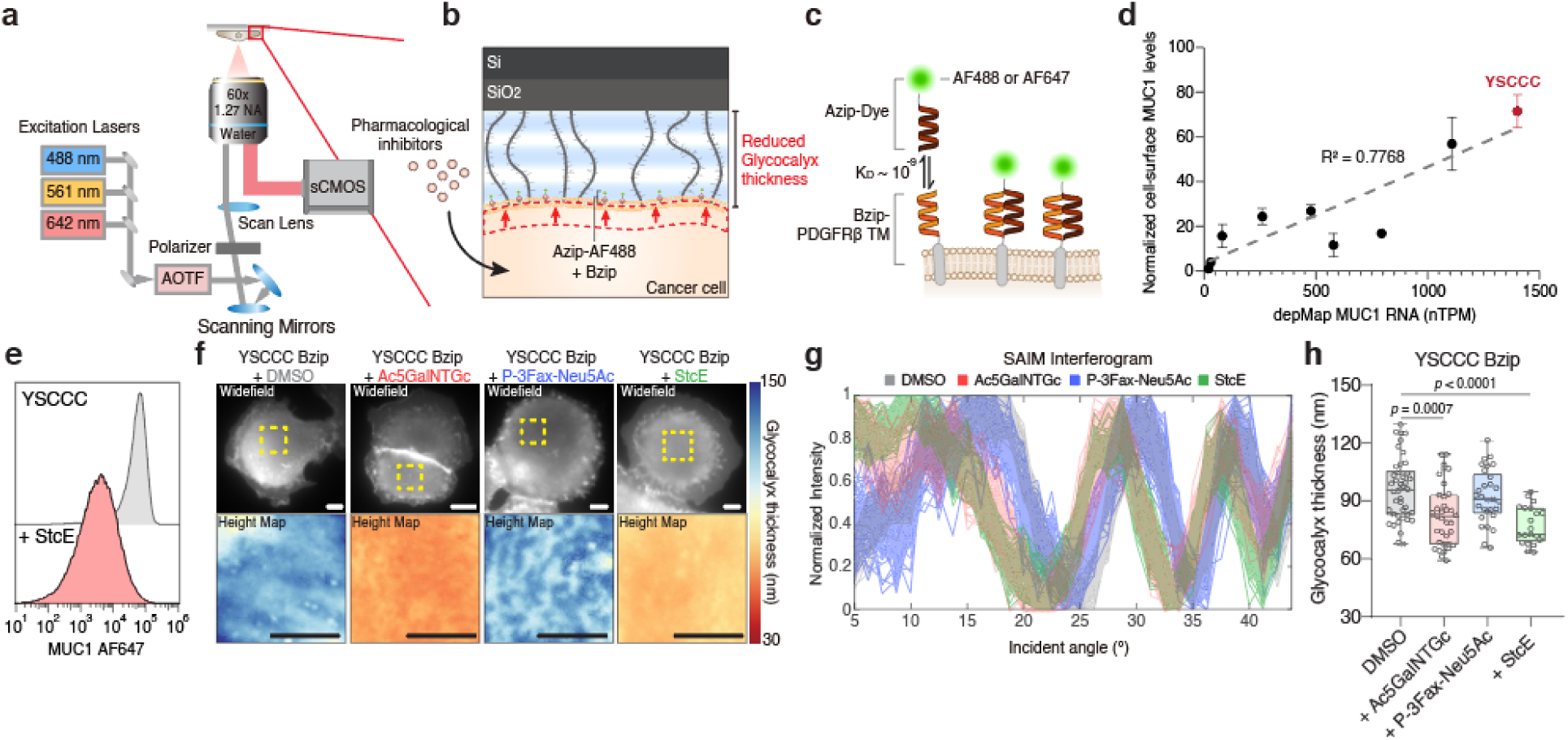
Membrane-referenced SAIM enables direct measurement of pharmacologically induced changes in cancer-cell glycocalyx thickness. **a**, Optical configuration of scanning angle interference microscopy (SAIM). Fluorescence intensity is recorded as a function of illumination angle to determine the axial position of membrane-associated fluorophores relative to the reflective silicon substrate. **b**, Schematic representation of membrane-referenced SAIM measurement of the cancer-cell glycocalyx. The plasma membrane is fluorescently labeled with AZip-AF488 bound to membrane-anchored BZip, enabling nanoscale measurement of glycocalyx thickness and its reduction following pharmacological perturbation. **c**, Schematic of the leucine zipper-based membrane-labeling strategy used for SAIM. Fluorophore-conjugated AZip binds membrane-anchored BZip fused to the PDGFRβ transmembrane (TM) domain. **d**, Relationship between MUC1 transcript abundance obtained from DepMap and normalized cell-surface MUC1 levels measured by flow cytometry across the indicated cancer cell lines. Data represent mean ± s.d. from at least two independent experiments. YSCCC cells are highlighted in red. The dashed line represents a linear regression.**e**, Representative flow-cytometry histograms of cell-surface MUC1 in YSCCC cells with or without StcE treatment. **f**, Representative wide-field fluorescence images and corresponding SAIM-derived glycocalyx height maps of BZip-expressing YSCCC cells treated with DMSO, Ac_5_GalNTGc, P-3F_AX_-Neu5Ac, or StcE and labeled with AZip-AF488. Scale bars, 10 μm. **g**, Representative pixel-wise SAIM interferograms from cells under the indicated treatment conditions. **h**, Quantification of glycocalyx thickness in BZip-expressing YSCCC cells treated with DMSO, Ac_5_GalNTGc, P-3F_AX_-Neu5Ac, or StcE. Boxes indicate the first and third quartiles, center lines indicate the median, and whiskers indicate the range. Each condition includes at least 21 cells from a representative experiment. Statistical significance was determined by one-way ANOVA followed by Tukey’s multiple-comparisons test.

We next tested whether AZip-BZip membrane labeling could provide a sufficiently stable reference for direct SAIM measurement of pharmacologically induced glycocalyx changes. BZip-expressing YSCCC cells were cultured on reflective silicon substrates and labeled with AZip-AF488 before SAIM acquisition. We first tested Ac_5_GalNTGc, P-3F_AX_-Neu5Ac, and StcE, which target *O*-glycosylation, sialylation, and cell-surface mucins, respectively. SAIM-derived height maps revealed the changes in glycocalyx physical thickness following each treatment compared with DMSO-treated cells. (**Fig. 3f**). Corresponding pixel-wise interferograms exhibited reproducible shifts across illumination angles, demonstrating that the membrane-referenced SAIM signal was sensitive to nanoscale changes induced by glycocalyx perturbation (**Fig. 3g**). Quantitative analysis showed that YSCCC cells had an average glycocalyx thickness of approximately 96 nm under control conditions. Ac_5_GalNTGc treatment reduced glycocalyx thickness to approximately 83 nm, corresponding to an average reduction of 13 nm, whereas StcE treatment reduced glycocalyx thickness to approximately 77 nm, a reduction of approximately 19 nm (**Fig. 3h**). In contrast, P-3F_AX_-Neu5Ac did not significantly alter glycocalyx thickness in YSCCC cells despite its established effects on cell-surface sialylation. Thus, changes in individual glycan features do not necessarily translate directly into predictable changes in the physical thickness of the glycocalyx.

To assess the broader applicability of the AZip-BZip labeling strategy, we performed SAIM measurements in multiple cancer cell lines, including KPL-1, SKBR3, and YSCCC, and obtained robust glycocalyx height maps across all three models (**Supplementary Fig. 3b-d**). Having established that SAIM could be applied across these models, we next directly compared the response of KPL-1 and YSCCC cells to the same pharmacological perturbations. In KPL-1 breast cancer cells, Ac_5_GalNTGc reduced the average glycocalyx thickness from approximately 68 nm to 47 nm, whereas P-3F_AX_-Neu5Ac produced a more pronounced reduction to approximately 28 nm (**Supplementary Fig. 3e,f**). This response differed markedly from that observed in YSCCC cells, in which P-3F_AX_-Neu5Ac had little effect on glycocalyx thickness. To determine whether this difference could be explained by the underlying glycosylation programs of the two cell lines, we compared transcriptome-inferred glycan biosynthetic pathways in YSCCC and KPL-1 cells using GlycoMaple^38^. Both cell lines expressed broad sets of enzymes supporting sialylated glycan biosynthesis, and the analysis did not reveal an obvious baseline difference that could explain the pronounced glycocalyx thinning in KPL-1 but not YSCCC cells following P-3F_AX_-Neu5Ac treatment (**Supplementary Figs. 7 and 8**). Thus, baseline transcriptional annotation of glycosylation pathways was insufficient to predict the physical response of the glycocalyx to sialylation inhibition. Together, these findings demonstrate that leucine zipper-based membrane labeling can be integrated with SAIM to resolve nanoscale changes in the physical structure of the cancer cell glycocalyx across different cell types and highlight the value of measuring glycocalyx thickness itself when evaluating pharmacological perturbations.

### SAIM-based pharmacological screening reveals diverse mechanisms of glycocalyx remodeling

Encouraged by these results, we next applied leucine zipper-based SAIM to screen an expanded panel of pharmacological compounds for their ability to reduce cancer-cell glycocalyx thickness (**Fig. 4a,b**). Because YSCCC cells form a particularly thick, MUC1-rich glycocalyx, we used this model as the primary platform for the initial thickness-based screen. The compound panel included inhibitors of glycan biosynthesis and processing, as well as metabolic pathways that can influence the production or modification of cell-surface glycoconjugates (**Fig. 4b**). Although many of these compounds have well-established biochemical targets, their effects on the physical thickness of the cancer-cell glycocalyx have not been systematically examined. We therefore quantified glycocalyx thickness in BZip-expressing YSCCC cells following treatment with each compound and compared these measurements with DMSO-treated controls. Most compounds reduced glycocalyx thickness, although the magnitude of the effect varied substantially across treatments (**Fig. 4c,d**).

**Figure 4.**
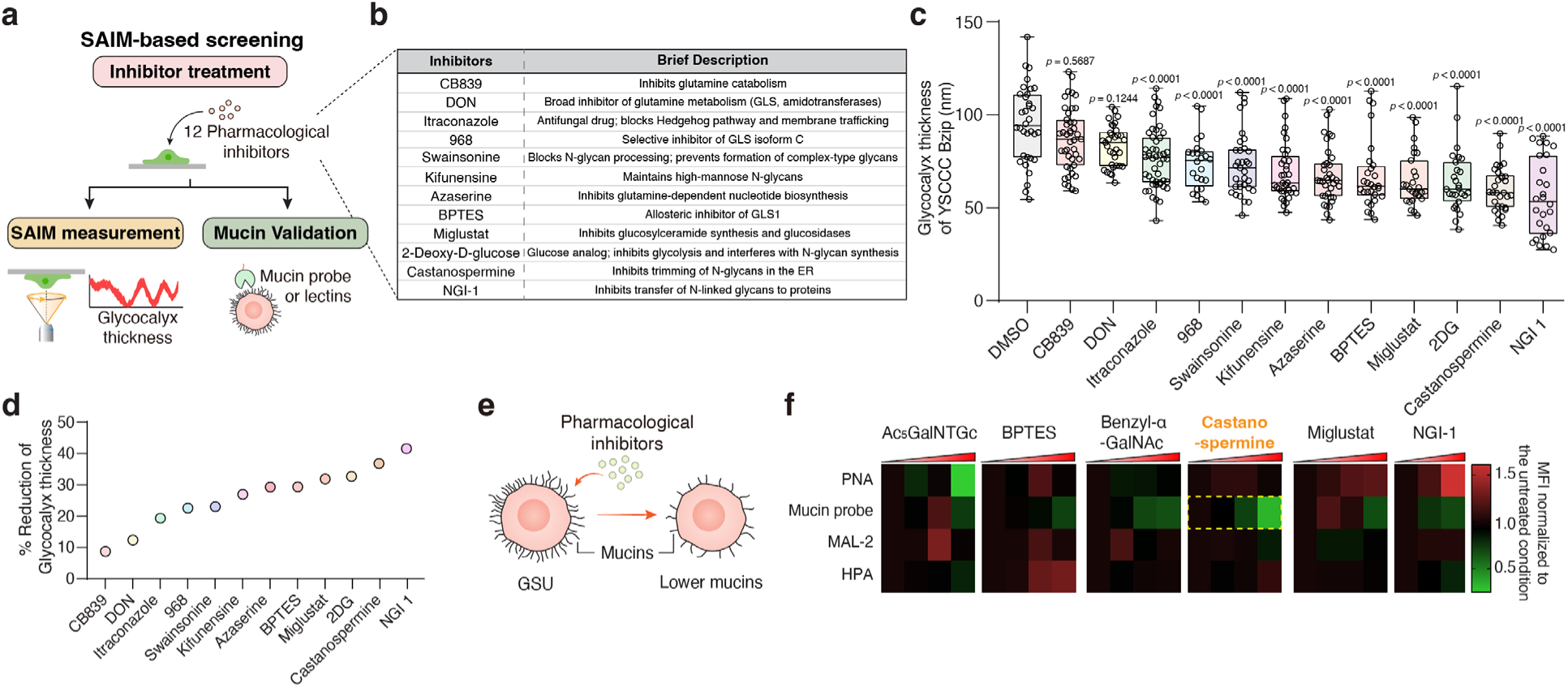
SAIM-based pharmacological screening and orthogonal mucin validation identify castanospermine as a lead glycocalyx-modulating compound. **a**, Schematic overview of the SAIM-based screening workflow. YSCCC cells were treated with a panel of pharmacological inhibitors and subsequently analyzed for glycocalyx thickness by SAIM. Candidate hits were further evaluated by orthogonal mucin/lectin-based validation to identify compounds that reproducibly remodel the cancer-cell glycocalyx. **b**, Summary table of the pharmacological inhibitors included in the screen, their primary molecular targets, and the glycan-processing steps they affect. **c**, Quantification of glycocalyx thickness in BZip-expressing YSCCC cells following treatment with the indicated inhibitors. Each point represents a single cell. Boxes indicate the first and third quartiles, center lines indicate the median, and whiskers indicate the range. Statistical significance was determined by one-way ANOVA followed by Tukey’s multiple-comparisons test. **d**, Ranked summary showing the percent reduction in glycocalyx thickness induced by each inhibitor relative to DMSO-treated controls. **e**, Schematic of orthogonal validation in GSU cells. Candidate compounds identified by SAIM screening were further assessed for their ability to alter cell-surface mucin features using mucin probes and lectins. **f**, Heat map summarizing changes in mucin-probe and lectin binding in GSU cells following treatment with the indicated inhibitors, normalized to untreated controls. Castanospermine emerged as a lead compound based on its reduction of glycocalyx thickness by SAIM together with reproducible changes in cell-surface probe binding.

The compounds that reduced YSCCC glycocalyx thickness spanned several mechanistically distinct classes. Although YSCCC cells were selected for their prominent mucin-rich glycocalyx, substantial structural changes were produced not only by perturbing mucin-associated glycosylation, but also by inhibiting protein *N*-glycosylation with NGI-1, early *N*-glycan processing with castanospermine, glycosphingolipid and ganglioside biosynthesis with miglustat, glutamine metabolism with BPTES and compound 968, and glucose metabolism with 2-deoxy-D-glucose (2DG) (**Fig. 4c,d**). NGI-1 produced the largest reduction in glycocalyx thickness, decreasing thickness by more than 40% relative to DMSO-treated cells, with castanospermine also among the strongest hits. Thus, even a mucin-rich glycocalyx could be remodeled through diverse glycan, lipid, and metabolic pathways, supporting glycocalyx thickness as an integrated cellular phenotype rather than a direct readout of any single biosynthetic pathway.

We next asked whether compound-induced glycocalyx thinning was associated with increased susceptibility to immune-cell attack. Several compounds that reduced glycocalyx thickness also increased NK-92-mediated cytotoxicity, although the magnitude of the effect varied among treatments (**Supplementary Fig. 6a**). Across the compounds tested, glycocalyx thickness showed a modest inverse relationship with NK-92-mediated cytotoxicity (Pearson r = -0.616; **Supplementary Fig. 6b**), consistent with a thicker glycocalyx providing greater protection from immune-cell engagement. However, compounds producing similar degrees of thinning did not necessarily produce equivalent changes in cytotoxicity, indicating that glycocalyx thickness captures an important physical component of the response but does not alone determine the functional outcome.

To further evaluate selected hits in a tumor context relevant to cellular immunotherapy, we turned to GSU gastric cancer cells, which display a mucin-rich cell surface that includes the MUC17 target antigen and serve as an established model for MUC17 CAR-T cells^39^. GSU cells were treated with selected compounds from the SAIM screen, and binding of PNA, MAL-II, HPA, and a mucin-directed probe was quantified by flow cytometry (**Fig. 4e,f** and **Supplementary Fig. 9**). The compounds produced distinct patterns of cell-surface remodeling. As expected, Ac_5_GalNTGc prominently reduced PNA binding, consistent with inhibition of mucin-type *O*-glycosylation, but produced comparatively limited changes in the other probes. Castanospermine produced the most pronounced reduction in mucin-probe binding, whereas miglustat and BPTES produced more modest reductions (**Fig. 4f).** Thus, compounds identified by their effects on glycocalyx thickness were accompanied by distinct changes in cell-surface glycan and mucin features in GSU cells.

Castanospermine emerged as a strong candidate for further study because it was among the most potent glycocalyx-thinning compounds in YSCCC cells and produced the most pronounced reduction in mucin-probe binding in GSU cells. Although NGI-1 produced the greatest reduction in glycocalyx thickness, it broadly inhibits protein *N*-glycosylation and can impair glycoprotein folding and induce ER stress^40^, raising concern that such effects could also compromise immune-effector cells during combination therapy. In contrast, castanospermine inhibits ER α-glucosidases involved in early *N*-glycan processing^41–43^ and has previously shown pharmacological activity following systemic and oral administration *in vivo*^44,45^, providing additional precedent for its further evaluation. We therefore selected castanospermine for deeper analysis of glycocalyx remodeling and its effects on MUC17 CAR-T-cell activity.

### Castanospermine-mediated glycocalyx remodeling enhances MUC17 CAR-T cell avidity and cytotoxicity

Having identified castanospermine as a prominent glycocalyx-thinning hit in YSCCC cells and observed reduced mucin-probe binding in GSU cells, we next characterized the transcriptional response to castanospermine in GSU cells before testing its effects on MUC17 CAR-T cell engagement and activity. Bulk RNA sequencing revealed a marked transcriptional response, with castanospermine-treated and control samples separating clearly by principal component analysis (**Fig. 5a**). Differential-expression analysis showed changes across multiple glycocalyx biosynthetic pathways, suggesting that the cellular effects of castanospermine may extend beyond its annotated role in early *N*-glycan processing (**Fig. 5b,c**). Among these changes, castanospermine decreased expression of MUC1 and MUC17, along with HAS2 and HAS3, which encode plasma-membrane hyaluronan synthases that generate extended hyaluronan chains (**Fig. 5b,c** and **Supplementary Fig. 10**). Consistent with a structural contribution of hyaluronan to the glycocalyx, doxycycline-inducible HAS3 expression progressively increased glycocalyx thickness with increasing doxycycline concentrations (**Supplementary Fig. 11a**). Together, these transcriptional changes suggested that castanospermine could alter multiple components of the glycocalyx and thereby modify the physical environment encountered by CAR-T cells at the tumor-cell surface. We therefore tested castanospermine in our established GSU-MUC17 CAR-T mode^39^.

**Figure 5.**
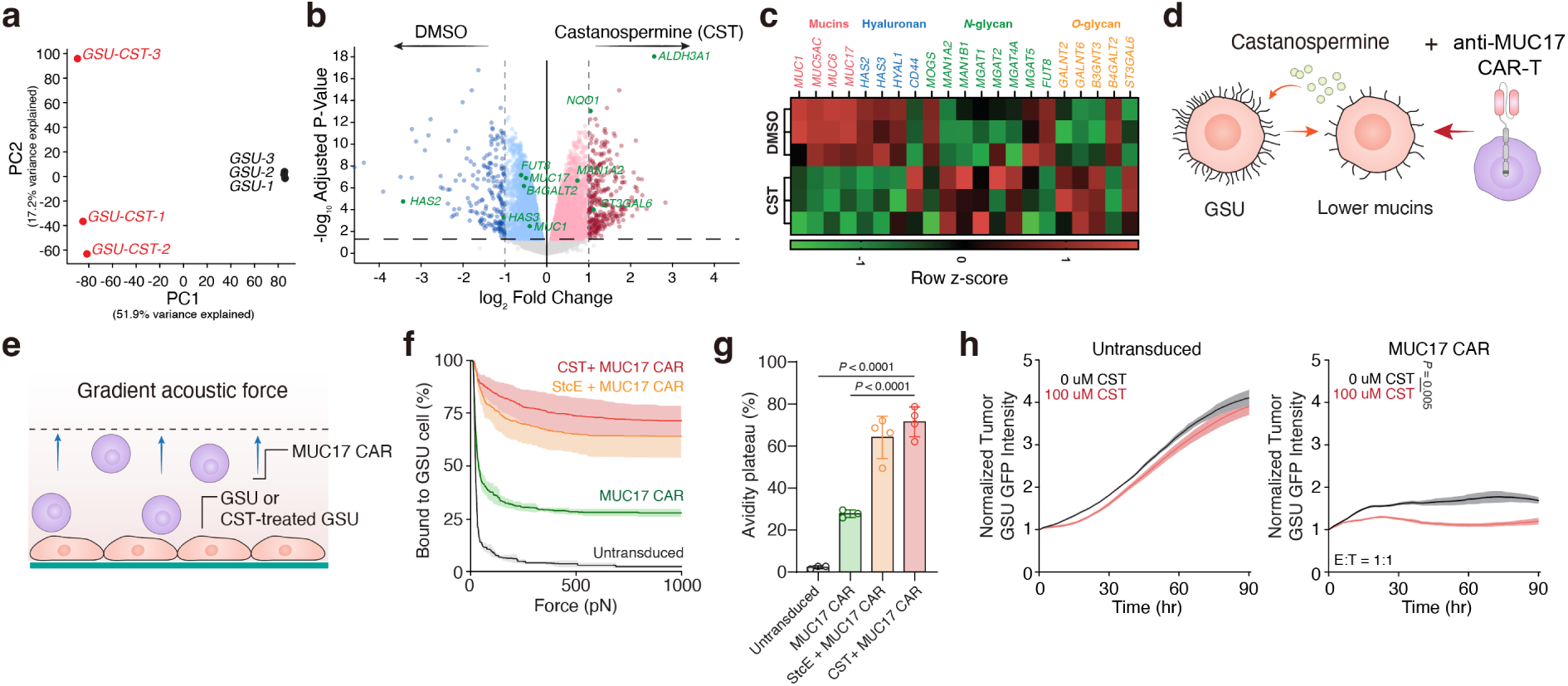
Castanospermine remodels the cancer cell glycocalyx and enhances MUC17 CAR-T cell avidity and cytotoxicity. **a**, Principal component analysis (PCA) of bulk RNA-sequencing profiles from GSU gastric cancer cells treated with DMSO or castanospermine (CST). Each point represents an independent biological replicate. The percentages of variance explained by PC1 and PC2 are indicated on the respective axes. **b**, Volcano plot showing differential gene expression in CST-treated versus DMSO-treated GSU cells. Genes enriched following CST treatment are shown to the right and genes enriched in DMSO-treated cells to the left. Selected genes associated with mucins, glycosaminoglycans and glycosylation pathways are highlighted and labeled. The dashed lines indicate the thresholds used to define differential expression. **c**, Heatmap of selected glycocalyx-associated genes in DMSO- and CST-treated GSU cells, grouped according to mucin, hyaluronan, *N*-glycan, and *O*-glycan-associated functions. Each row represents an independent biological replicate. Gene expression values were normalized as counts per million (CPM), log_2_-transformed, and displayed as row-wise z-scores. **d**, Schematic illustrating CST-mediated remodeling of the GSU cancer cell glycocalyx followed by targeting with MUC17 CAR-T cells. **e**, Schematic of the gradient acoustic force assay used to quantify cell-cell avidity between MUC17 CAR-T cells and untreated or glycocalyx-modulated GSU target cells. **f**, CAR-T cell avidity profiles measured under increasing acoustic force for untransduced T cells or MUC17 CAR-T cells interacting with untreated, StcE-treated, or castanospermine-treated GSU cells. The percentage of CAR-T cells remaining bound to GSU target cells is plotted as a function of applied force. **g**, Quantification of the avidity plateau derived from the force curves shown in **f**. Results are mean ± s.e.m. for force-ramp measurements and mean ± s.d. for avidity-plateau measurements from n = 3 independent measurements. **h**, Real-time cytotoxicity assay of GSU target cells co-cultured with untransduced T cells or MUC17 CAR-T cells in the presence or absence of 100 μM castanospermine. Cancer cell growth was monitored by normalized GSU GFP fluorescence intensity over time. MUC17 CAR-T cells were co-cultured with GSU cells at an E:T ratio of 1:1. Results are mean ± s.d. of n = 3 independent measurements. Statistical analysis in **g** was performed using one-way ANOVA with Tukey’s multiple-comparisons test. In **h,** statistical analysis was performed by two-way ANOVA with correction for multiple comparisons.

Before assessing CAR-T cell engagement, we first determined whether castanospermine altered the abundance of the MUC17 target antigen on GSU cells. Consistent with the transcriptional decrease in *MUC17*, cell-surface MUC17 antibody binding showed a modest decrease following castanospermine treatment (**Supplementary Fig. 11b**). We next asked whether castanospermine could nevertheless enhance physical engagement between GSU cells and MUC17 CAR-T cells through remodeling of the tumor-cell surface (**Fig. 5d**). Cell-cell avidity was measured using a gradient acoustic force assay, in which increasing acoustic force was applied to detach T cells bound to adherent GSU target cells (**Fig. 5e**). MUC17 CAR-T cells displayed substantially greater avidity toward GSU cells than untransduced T cells, and enzymatic removal of cell-surface mucins with StcE further increased CAR-T cell binding (**Fig. 5f,g**). Importantly, castanospermine treatment produced a comparable enhancement of cell-cell avidity, increasing the avidity plateau by approximately 43.7 percentage points relative to untreated GSU cells (**Fig. 5f,g**). Thus, despite modestly reduced MUC17 abundance, pharmacological remodeling of the tumor-cell glycocalyx markedly enhanced physical engagement between MUC17 CAR-T cells and their targets.

We next determined whether this increased avidity translated into improved tumor control. GFP-expressing GSU cells were co-cultured with either untransduced T cells or MUC17 CAR-T cells in the presence or absence of 100 μM castanospermine, and tumor growth was monitored in real time. Castanospermine produced only a modest effect on GSU growth in cultures containing untransduced T cells (**Fig. 5h**). In contrast, the addition of castanospermine to MUC17 CAR-T cell cultures significantly improved tumor control compared with MUC17 CAR-T cells alone (**Fig. 5h**). Together with the avidity measurements, these findings support a model in which castanospermine enhances MUC17 CAR-T activity by remodeling the physical tumor-cell surface and improving CAR-T-target engagement, rather than by increasing antigen abundance or simply suppressing cancer-cell growth.

We further evaluated several additional compounds identified in the SAIM screen to determine whether glycocalyx thinning alone was sufficient to enhance MUC17 CAR-T activity (**Supplementary Fig. 12**). NGI-1, which produced the greatest reduction in glycocalyx thickness in the initial screen, also reduced GSU growth at 1.5 μM but did not produce a corresponding enhancement of MUC17 tumor control. Similarly, benzyl-α-GalNAc and miglustat produced little change in either cancer cell growth or CAR-T cell-mediated tumor control. BPTES appeared to improve tumor control in the presence of MUC17 CAR-T cells, but it also reduced GSU growth in cultures containing untransduced T cells, indicating a substantial CAR-independent component. In contrast, castanospermine enhanced tumor control preferentially in the MUC17 CAR-T condition. These results identify castanospermine as a glycocalyx-modulating compound that improves MUC17 CAR-T cell engagement and activity, while highlighting the value of combining glycocalyx-thickness measurements with functional testing to identify effective immunotherapy combinations.

## Discussion

The physical dimensions of the cancer-cell glycocalyx are a modifiable determinant of immune-cell killing. Here, we establish glycocalyx thickness as a druggable physical phenotype that can be measured directly and used to guide the selection of compounds for combination with cellular immunotherapy. Pharmacological disruption of mucin-type *O*-glycosylation reduced glycocalyx thickness and enhanced NK- and CAR-NK-mediated cytotoxicity. To extend this principle to compound screening, we developed a stable extracellular plasma-membrane reference based on a complementary leucine-zipper pair and integrated it with SAIM, enabling nanoscale measurements across cancer cell types. A focused screen, followed by orthogonal mucin and lectin profiling, identified castanospermine as a candidate glycocalyx-modulating agent that increased MUC17 CAR-T cell avidity and improved tumor control in gastric cancer co-cultures. Together, these findings connect a measurable physical property of the cancer-cell surface to a pharmacological strategy for improving cellular immunotherapy.

A defining feature of the glycocalyx thickness phenotype is the small physical scale over which it operates. For instance, *O*-glycosylation inhibition reduced glycocalyx thickness by approximately 14 nm, and in the CD19 CAR-NK model the same perturbation increased cytotoxicity 3.02-fold relative to CAR-NK cells against untreated targets. Previous studies demonstrated that enzymatic mucin removal can improve immune-cell attack^2,10,11^, while disruption of tumor *N*-glycosylation with 2-deoxy-D-glucose enhanced CAR-T activity in several solid-tumor models^13^. The present results place these functional effects on a directly measured physical scale. Importantly, enhanced killing followed partial thinning of the glycocalyx, suggesting that therapeutic benefit need not require complete removal of a mucin or glycan class.

Direct measurement of glycocalyx thickness is particularly informative because the structural response to glycosylation inhibitors depends on cellular context. KPL-1 cells were particularly sensitive to sialylation inhibition by P-3F_AX_-Neu5Ac^15,16^, whereas the same treatment produced little change in YSCCC glycocalyx thickness. YSCCC cells instead showed a clear response to perturbation of mucin-associated glycosylation. Although transcriptome-based analysis with GlycoMaple^38^ revealed differences in the baseline glycosylation programs of these models, those profiles did not readily explain their distinct structural responses. The physical structure of the glycocalyx emerges from enzyme activity, substrate availability, glycan processing, and the abundance and organization of cell-surface glycopolymers in addition to transcript levels. Therefore, thickness provides an integrated readout of the physical structure produced by these processes. Pairing structural measurements with molecular profiling can distinguish what a compound changes biochemically from whether those changes alter the physical barrier encountered by an immune cell.

Castanospermine further illustrates why the structural consequences of a biochemical perturbation can extend beyond its annotated target. Its established biochemical activity is inhibition of ER α-glucosidases involved in early *N*-glycan processing.^41,42^ Yet in treated GSU cells, the transcriptional response extended across multiple classes of glycocalyx components and biosynthetic pathways. Expression of *MUC1* and *MUC17* decreased alongside *HAS2* and *HAS3*, whereas genes involved in *N*- and *O*-glycosylation changed in both directions. Thus, inhibition of a single *N*-glycan-processing step was accompanied by broader changes in pathways that could act in opposing directions on the physical structure of the glycocalyx. The net surface phenotype therefore cannot be inferred from the direction of the inhibited step alone. Although glycocalyx thickness was not measured directly in castanospermine-treated GSU cells, the reduction in mucin- probe binding together with the transcriptional response supports broader remodeling of the cell surface. These observations suggest that glycosylation inhibitors can reshape the glycocalyx through secondary transcriptional and biosynthetic responses in addition to their direct effects on glycan processing.

The identification of metabolic inhibitors, including BPTES and compound 968, as glycocalyx-thinning agents extends this principle beyond compounds conventionally classified as glycosylation inhibitors. Nutrient metabolism and glycan biosynthesis are closely coupled through the hexosamine biosynthetic pathway, and prior work in T cells has shown that glycolysis and glutaminolysis can regulate *N*-glycosylation by controlling metabolite supply.^46^ In our screen, both BPTES and 968 reduced glycocalyx thickness, revealing a physical cell-surface response to glutaminase inhibition. Because glutaminase inhibition redistributes glutamine utilization rather than directly targeting a glycosyltransferase, the route from metabolic perturbation to glycocalyx thinning is unlikely to be captured by the annotated target alone. The concordant response to two chemically distinct inhibitors suggests that metabolic interventions can alter the physical structure of the glycocalyx in addition to their effects on cell growth. More broadly, thickness-based screening can capture such integrated surface responses without requiring prior knowledge of every intermediate pathway through which a compound acts.

A further implication for CAR-T therapy is that antigen abundance and physical accessibility are separable determinants of target engagement. Target-antigen density and CAR design are well-established determinants of CAR-T activity.^47^ Yet classical physical models of cell adhesion describe productive binding as a competition between specific receptor-ligand bonds and nonspecific surface repulsion, including steric forces generated by macromolecular glycocalyx layers.^48,49^ This framework is particularly relevant to immune synapses, which require close apposition of effector and target membranes. Here, castanospermine increased the avidity plateau of MUC17 CAR-T cells for GSU targets by approximately 43.7 percentage points, to a level comparable to enzymatic mucin removal with StcE, despite only a modest decrease in MUC17 antibody binding. Notably, GSU cells were pretreated and washed before the avidity assay, arguing that the increased engagement resulted from changes to the tumor-cell surface rather than acute drug exposure of the T cells. Although direct thickness measurements were obtained in YSCCC rather than GSU cells, the combination of reduced mucin-probe binding and increased avidity in GSU cells supports a model in which glycocalyx remodeling improves physical access to the target-cell surface. Thus, CAR-T engagement can be improved without increasing antigen abundance or changing CAR design, providing a complementary strategy to efforts that optimize antigen expression or receptor properties.

The comparison of screening hits also illustrates why glycocalyx thickness is most useful as an entry point for compound selection rather than a stand-alone predictor of therapeutic benefit. Agents that remodel the tumor-cell surface can also act on effector cells, making functional triage an important second step. NGI-1 produced the largest reduction in YSCCC glycocalyx thickness but did not enhance MUC17 CAR-T-mediated tumor control in GSU co-cultures. As an inhibitor of the oligosaccharyltransferase complex^40^, NGI-1 broadly suppresses protein *N*-glycosylation, a process that is also important for T-cell function^46^. BPTES improved tumor control in the CAR-T condition but also reduced GSU growth in cultures containing untransduced T cells, indicating a substantial CAR-independent component. Castanospermine, by contrast, improved tumor control preferentially in the condition of MUC17 CAR-T cell co-culture. Studies *in vivo* can now determine how dosing and delivery influence the balance among tumor-surface remodeling, effector-cell fitness, and effects on normal tissues.

Together, these findings establish a drug-discovery strategy organized around a physical property of the cancer-cell surface. Glycocalyx thickness provides a measurable integrated output of diverse biosynthetic and regulatory processes, while molecular profiling and functional assays identify forms of remodeling that improve immune-cell engagement. Although the focused panel used here was small enough to permit parallel or independent functional testing, it allowed us to test whether glycocalyx thickness could be used to guide compound selection. The value of this approach should increase as chemical space for compound discovery expands, where exhaustive functional testing becomes less practical and annotated molecular targets provide limited guidance about the glycocalyx physical structure. Castanospermine provides proof of concept for using this strategy to identify a small-molecule partner for MUC17 CAR-T cells. More broadly, the physical environment in which a tumor antigen is presented can be pharmacologically modified alongside the receptors that recognize it, making the physical structure of the glycocalyx an actionable design parameter for cellular immunotherapy.

## METHODS

### Cell culture

MCF10A cells were maintained in DMEM/F12 medium (Thermo Fisher Scientific) supplemented with 5% horse serum (Thermo Fisher Scientific), 20 ng/ml EGF (PeproTech), 10 μg/ml insulin (Sigma), 500 ng/ml hydrocortisone (Sigma), 100 ng/ml cholera toxin (Sigma), and 1x penicillin/streptomycin (Thermo Fisher Scientific) at 37°C with 5% CO_2_. HEK293T cells (gift from Valerie Weaver) were cultured in DMEM high glucose media (Thermo Fisher Scientific) containing 10% fetal bovine serum (Thermo Fisher Scientific) and 1x penicillin/streptomycin at 37°C in 5% CO_2_. YSCCC (RIKEN BRC; RCB1549), OE19 (DSMZ; ACC 700), ZR-75-1 (ATCC; CRL-1500), T47D (ATCC; HTB-133), SKBR3 (gift from Dr. Jan Lammerding), KPL-1 (DSMZ; ACC 317), and Capan-2 (ATCC; HTB-80), and GSU cells were cultured in RPMI 1640 media (Thermo Fisher Scientific) supplemented with 10% fetal bovine serum and 1x penicillin/streptomycin at 37°C in 5% CO_2_. SNU-C2A cells (ATCC; CCL-250.1) were cultured in DMEM/F12 medium (Thermo Fisher Scientific) supplemented with 10% fetal bovine serum, 0.02 mg/mL insulin, 0.01 mg/mL transferrin, 25 nM sodium selenite, 50 nM hydrocortisone, 1 ng/mL EGF, 0.01 mM ethanolamine, 0.01 mM phosphorylethanolamine, 100 pM triiodothyronine, 0.5% (w/v) bovine serum albumin, 10 mM HEPES, 0.5 mM sodium pyruvate, 2 mM L-glutamine, and 1x penicillin/streptomycin at 37°C with 5% CO_2_. NK-92 cells (ATCC; CRL-2407) were cultured in ɑ-MEM without ribonucleosides media (Thermo Fisher Scientific) supplemented with 12.5% fetal bovine serum, 12.5% horse serum, 0.2 mM Myo-inositol (Sigma Aldrich), 0.1 mM 2-mercaptoethanol (Thermo Fisher Scientific), 0.02 mM folic acid (Millipore Sigma), 100 U/ml recombinant human IL-2 (Pepro Tech), and 1x penicillin/streptomycin at 37°C in 5% CO_2._

### Doxycycline-inducible MUC1 and *HAS3* cell model

The doxycycline-inducible MUC1-GFP MCF10A model or *HAS3* MCF10A was generated as previously described^2,50^. Briefly, MUC1-GFP containing 42 tandem repeats was stably expressed under the control of a tetracycline-inducible promoter, and the clonally expanded 1E7 line was selected for its titratable MUC1 surface expression. For cell-surface MUC1 density experiments, cells were treated with the indicated concentrations of doxycycline for 24 h before subsequent enzyme treatment, flow cytometric analysis or cytotoxicity assays.

### Generation of BZip-expressing cell lines

BZip-expressing cell lines were generated by lentiviral transduction with constructs encoding BZip fused to a PDGFRβ transmembrane (TM) domain. BZip plasmids were synthesized by Twist Bioscience. Following transduction, mScarlet-positive cells were isolated using a Sony MA900 cell sorter to establish stable BZip-expressing populations. BZip-expressing Capan-2, KPL-1, T47D, SKBR3, ZR-75-1, and YSCCC cell lines were generated using this approach.

### Immunostaining and live cell imaging

Target cells were plated in 35-mm glass-bottom dishes and cultured for 24 hours. For standard conditions, cells were labeled with 1 µM AZip-based dyes (AZip-sfGFP, AZip-AF488, or AZip-AF647) in phenol red-free culture media for 10 minutes at 37°C. For cold conditions, cells were labeled with 1 µM AZip-based dyes in 0.5% BSA PBS for 1 hour at 4°C. For 48-hour live cell imaging experiments, cells were labeled according to manufacturers’ recommended buffer compositions and concentrations for 10 minutes at 37°C. Before imaging, cells were washed twice with either cold PBS or phenol red-free culture media. All samples were imaged at 0, 4, 24, and 48 hours at 37°C under 5% CO_2_. For live cell imaging of 1E7 cells in response to inhibitors, cells were first induced with various concentrations of doxycycline for 24 hours, then treated with either DMSO or 100 μM Ac_5_GalNTGc for 48 hours. All imaging was performed using an LSM 800 confocal microscope with either a x20 air objective (NA 0.8) or x63 water objective (NA 1.2) (ZEISS). ImageJ was used for image analysis.

### CAR-T cell production

Human CAR-T cells were generated from de-identified leukapheresis products obtained from healthy donors through the MGH Blood Bank under an Institutional Review Board-exempt protocol, as previously described^39^. T cells were isolated using the RosetteSep T Cell Enrichment Cocktail (STEMCELL Technologies) and activated with CD3/CD28 Dynabeads (Life Technologies) at a 1:3 T cell-to-bead ratio. Cells were maintained in RPMI 1640 containing GlutaMAX and HEPES, supplemented with 10% fetal bovine serum, penicillin/streptomycin, and 20 IU/mL recombinant human IL-2 (PeproTech). At 24 h after activation, cells were transduced with CAR-encoding lentivirus at a multiplicity of infection of 5-10 and expanded at approximately 0.5-1 × 10^6^ cells/mL. Dynabeads were removed on day 6, and CAR expression was evaluated by flow cytometry before cryopreservation. Untransduced T cells generated from the same donors were used as controls. Before functional assays, CAR-T and untransduced T cells were thawed and rested for 18-24 h in IL-2-containing RPMI 1640 medium.

### Flow cytometry

To measure cell surface MUC1 expression in cancer cell lines, cells were plated, grown for at least 48 hours, and detached using trypsin. Anti-MUC1 antibody clone HMPV (555925, BD Biosciences) was diluted 1:200 in 0.5% w/v BSA in 1x PBS and incubated with the cells at 4°C for 1 hour. Secondary labelling was with Alexa Fluor 647 conjugated goat anti-mouse, diluted 1:200 in 0.5% w/v BSA in PBS and incubated with cells at 4°C for 1 hour. In each experiment, SKBR3 cells were included as a standard for comparison. Median fluorescence intensity was calculated in FlowJo and values were normalized to the median fluorescence intensity of SKBR3 cells in each experiment. For lectin or mucin staining, detached target cancer cells were incubated with PNA-CF640R (Biotium), biotinylated MAL-II (B-1265-1, Vector Laboratories), HPA-AF647 (Thermo Fisher Scientific), or mucin probe (SAE0212, Milipore Sigma) at 4°C for 1 hour. Lectins were diluted at 1:200 in 0.5% BSA PBS and incubated with cells at 4°C for 1 hour. For MUC17 staining, cells were incubated with APC-conjugated anti-MUC17 antibody (clone MQ155; BioLegend) at a 1:100 dilution in 0.5% BSA PBS and incubated with cells at 4°C for 1 hour. The Attune NxT flow cytometry (Thermo Fisher Scientific) was used for analysis.

### Sortase-mediated labeling reaction

Fluorescent AZip probes were generated by sortase-mediated labeling as previously described^51^. Briefly, 100 μM AZip-LPETG-sfGFP was incubated in reaction buffer (50 mM Tris-HCl, pH 7.5, 150 mM NaCl, 10 mM CaCl2) containing 20 μM sortase and 10 mM triglycine (Gly_3_)-conjugated AF488 or AF647. After 1-3 hours of incubation at room temperature, the reaction products were purified using HisPur Ni-NTA Resin to remove unreacted His6-tagged AZip dye and His6-tagged Sortase in elution buffer (20 mM HEPES, 500 mM NaCl, 10 mM imidazole, pH 7.5). The column flow-through was then buffer-exchanged to PBS to remove unbound dye.

### Treatment of pharmacological inhibitors

For pharmacological inhibitor treatment on YSCCC or GSU cells, cells were treated with the given pharmacological inhibitor or DMSO vehicle control diluted in RPMI 1640 media (Thermo Fisher Scientific) supplemented with 10% fetal bovine serum and 1x penicillin/streptomycin at 37°C in 5% CO_2_ at the following concentrations. Miglustat (MedChemExpress HY-17020), Kifunesine (MedChemExpress HY-19332), Swainsonine (Tocris Bioscience 3208), NGI-1 (MedChemExpress HY-117383), DON (MedChemExpress HY-108357), were applied at 50 µM for 48 hours. Azaserine (MedChemExpress HY-B0919) was applied at 50 µM for 12 hours. Castanospermine (MedChemExpress HY-N2022), was applied at indicated concentration for 24-72 hours. 2-deoxy-glucose (2DG; MedChemExpress HY-13966) was applied at 5 mM for 24 hours. Itraconazole (Selleck Chemicals) was applied at 2.5 µM for 48 hours. BPTES (kind gift from the Cerione Lab) was applied at 50 µM for 24 hours, CB-839 (gift from the Cerione Lab) was applied at 1 µM for 24 hours, and glutaminase inhibitor 968 (gift from the Cerione Lab) was applied at 50 µM for 48 hours. Ac_5_GalNTGc was used at either 60 or 100 μM depending on the experiment, as specified in the corresponding figure legends.

### NK-92 cell-mediated cytotoxicity assays

NK-92 cell-mediated cytotoxicity was measured by flow cytometry as previously described^2^. Briefly, target cells were labeled with 10 μM CellTracker Green CMFDA (Invitrogen) for 15 min and washed twice before co-culture. Labeled target cells (4 × 10^4^) were mixed with NK-92 cells at the indicated effector-to-target ratios in 200 μL of target-cell growth medium without IL-2 and incubated for 4 h at 37°C with 5% CO_2_. Cells were then stained with 20 μg/mL propidium iodide (Sigma), and target-cell death was quantified by flow cytometry after gating on CellTracker Green-positive cells. Cytotoxicity was calculated relative to spontaneous target-cell death as previously described^2^.

### Cell avidity measurement by acoustic force

Cell avidity was measured using the z-Movi Cell Avidity Analyzer (LUMICKS) as previously described^39,52^. Briefly, untreated or inhibitor-treated GSU cells were washed and seeded onto poly-L-lysine-coated z-Movi chips at 6 × 10^7^ cells/mL for 2 h. CAR-positive T cells were enriched by PE bead-based sorting using PE-conjugated (G4S)_3_ antibody (Cell signaling Technology) before the assay and labeled with CellTrace Far Red (Thermo Fisher Scientific). MUC17 CAR-T cells or donor-matched untransduced T cells were allowed to bind to GSU cells for 5 min, followed by application of a gradually increasing acoustic force of up to 1,000 pN. The percentage of T cells remaining bound as a function of applied force was quantified using Ocean software (version 1.5.5; LUMICKS). Each condition was measured in at least three technical runs.

### Scanning angle interference microscopy (SAIM)

SAIM imaging and analysis were performed as previously described^2,22^. Briefly, silicon wafers containing an approximately 2,000-nm thermal oxide layer (Addison Engineering) were diced into 7 x 7 mm chips, and the oxide thickness of each chip was measured using a FilMetrics F50-EXR. Chips were functionalized with 4% (v/v) (3-mercaptopropyl)trimethoxysilane, followed by 4 mM 4-maleimidobutyric acid N-hydroxysuccinimide ester and 50 µg/mL Alexa Fluor 647-conjugated human plasma fibronectin as previously described^23^. Cells were seeded onto fibronectin-coated chips at 0.5-1 × 10^5^ cells/cm² and cultured for 24 h. For AZip-based SAIM measurements, cells were labeled with AZip-AF488 in phenol red-free medium for 10 min at 37°C and washed before imaging. For 1E7 cells, MUC1 expression was induced with the indicated concentrations of doxycycline for 24 h, followed by labeling with MemGlow 560 (MG02-2; Cytoskeleton) for 10 min at 37°C. Cell-seeded chips were inverted onto 35-mm glass-bottom dishes and imaged at 37°C. SAIM was performed using a custom circle-scanning microscope as previously described^2^. Images were acquired at 32 incidence angles ranging from 5° to 43.75°, and fluorescence intensity profiles were fit pixelwise to the established SAIM optical model using nonlinear least-squares fitting. Glycocalyx thickness was determined from the axial separation between the fluorescent plasma-membrane reference and the corresponding Alexa Fluor 647-labeled fibronectin substrate. Average thickness was quantified within 200 x 200-pixel regions of individual cells.

### Preparation of recombinant StcE mucinases

Recombinant StcE mucinases were produced and purified as previously described^2^. Briefly, proteins were expressed in NiCo21 (DE3) E. coli (NEB) and induced with 0.5 mM IPTG overnight at 24°C. Bacterial pellets were lysed by sonication, and recombinant proteins were purified by immobilized metal affinity chromatography using a HisTrap HP column (Cytiva), followed by size-exclusion chromatography on a HiPrep 26/60 Sephacryl S-200 HR column (Cytiva). Purified proteins were concentrated using 30-kDa MWCO Amicon Ultra centrifugal filters (MilliporeSigma) and stored in 20 mM HEPES and 150 mM NaCl (pH 7.5).

### Assessment of cell-substrate adhesion

YSCCC cells were plated at 80,000 cells per well in a 24-well plate (Cellstar, #662160) in RPMI 1640 media with 10% fetal bovine serum and 1% penicillin/streptomycin. Where specified, StcE or SmE mucinases were added to a final concentration of 1 nM. Cells were allowed to adhere to the plates for 18 hours in 37°C and 5% CO_2_. Phase contrast images were then acquired on a BZ-X810 fluorescence box microscope (Keyence) using a 20x (NA: 0.45) air objective. Cells were manually segmented and counted in ImageJ to calculate circularity and fraction of rounded cells (cells with a Circularity > 0.9). Rounded cell fraction was calculated as (# rounded cells)/(# rounded cells + # spread cells) in each field of view.

### Real-time cytotoxicity assays

CBG-eGFP-expressing GSU cells were seeded in flat-bottom 96-well plates (Corning) and allowed to adhere for at least 2 h at 37°C with 5% CO_2_. CAR-T cells or donor-matched untransduced T cells were then added at the indicated E:T ratios. Where indicated, pharmacological inhibitors were added at the start of co-culture. Plates were maintained at 37°C for up to 6 days and imaged every 2 h using the IncuCyte Live-Cell Analysis System. Tumor-cell growth and CAR-T cell-mediated cytotoxicity were quantified based on the total GFP-positive area using IncuCyte image-analysis software.

### RNA sequencing and analysis

Total RNA was isolated from three independent DMSO-treated and three castanospermine-treated GSU samples and submitted to Plasmidsaurus for library preparation, RNA sequencing, alignment, and gene-level quantification. Differential-expression and downstream analyses were performed using the processed gene-expression data provided by Plasmidsaurus. For heatmap visualization, expression values were transformed as log2(CPM + 1) and normalized by row-wise z-score across samples.

### Assessment of multicellular compaction

YSCCC cells were detached with 0.25% trypsin. Cells were seeded at 1,000 cells per well in 150 uL 96-well ultra-low adhesion plates (Corning #7007) in the presence of 0.05% methylcellulose (M6385, Millipore Sigma). Where specified, StcE was added to a final concentration of 10 nM. The plate was then gently centrifuged at 150xg for 3 minutes to ensure that all cells collected at the bottom of each well. Cells were then allowed to form aggregates for 24 hours at 37°C and 5% CO_2_. Cell aggregates were then imaged on an LSM 800 confocal microscope using the brightfield mode with a 10x (NA: 0.3 Air) objective. Aggregates were then manually segmented in ImageJ to calculate overall aggregate area.

### Statistical analysis

Unless otherwise indicated, results are presented as the mean and standard deviation (s.d.) of at least three replicates per condition using GraphPad Prism 9. Statistical differences were determined using a two-tailed unpaired t-test for two group comparisons, one-way ANOVA with multiplicity-adjusted p values from Tukey’s multiple comparisons test, and two-way ANOVA with correction for multiple comparisons. Statistical significance was determined using two-tailed unpaired t-tests, one-way ANOVA, and two-way ANOVA, with p < 0.05 considered significant across all analyses.

## Supporting information

Park Supplementary Information 2026

## Data availability

All data shown in this manuscript are provided as source data files. All additional data and materials that can be shared will be released using a material transfer agreement. Please contact Matthew J. Paszek and Sangwoo Park.

## Acknowledgments

SAIM instrument development was supported by the Kavli Institute at Cornell for Nanoscale Science. We thank Xiaolei Su for providing the CD19 CAR cDNA. This investigation was supported by the National Cancer Institute CA276398 (M.J.P.), CA238268 (M.V.M.), and CA303346 (S.P.), and Cornell IGNITE (G44TRAN, INOVA). Work was performed at the Cornell Nanoscale Facility (NSF NNCI-2025233), Biotechnology Resource Center (RRID:SCR_021740), and Imaging Facility (RRID:SCR_021741) with NYSTEM (C029155) and NIH (S10OD018516) funding for the Zeiss LSM880.

## Author Contributions

S.P. and M.J.P. designed the project. S.P. conducted the SAIM measurements and analysis. S.P. and J.H.P. conducted and analyzed all cytotoxicity assay and flow cytometry. S.P., J.H.P., and M.J.P. wrote the manuscript with feedback from all authors.

