## Supplementary material for "Cancer cell glycocalyx thickness is a druggable physical barrier to cellular immunotherapy": Park Supplementary Information 2026

**This Supplementary Information file contains:**

**- Supplementary Figures 1-11**

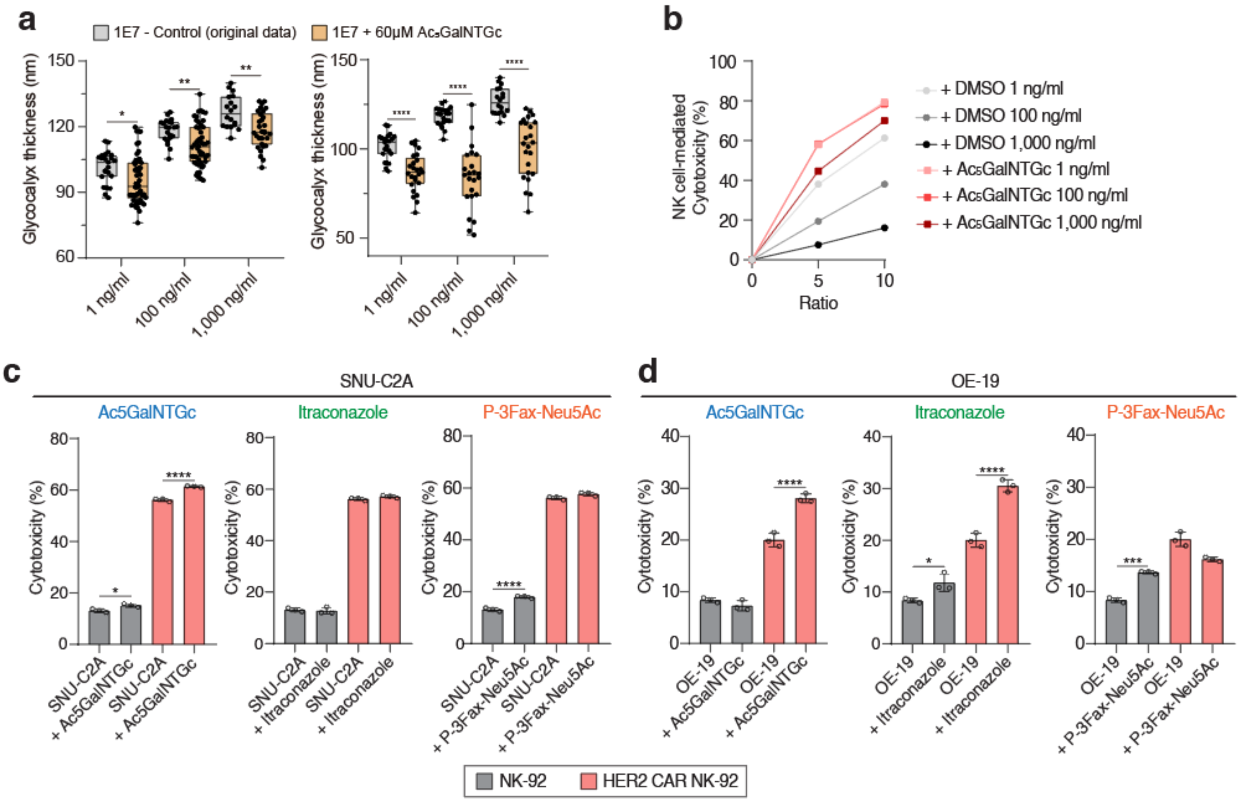


**Supplementary Figure 1. Glycocalyx modulation enhances NK- and CAR-NK-cell cytotoxicity across additional cancer cell lines. a,** Quantification of glycocalyx thickness in the doxycycline-inducible MUC1 1E7 clone cultured with the indicated concentrations of doxycycline and treated with DMSO or 60 µM Ac$_5$GalNTGc. Data are from an independent experiment replicating the analysis shown in **Fig. 1f**. Boxes indicate the first and third quartiles, center lines indicate the median, and whiskers indicate the range. **b**, NK-92 cell-mediated cytotoxicity against 1E7 target cells cultured with the indicated doxycycline concentrations and treated with DMSO or 60 µM Ac$_5$GalNTGc. Cytotoxicity was measured after 4 h of co-culture at the indicated E:T ratios. Cells were pretreated with doxycycline for 24 h, and inhibitor treatment was performed for 48 h before the cytotoxicity assay. c, Cytotoxicity of unmodified NK-92 cells or HER2 CAR-expressing NK-92 cells against SNU-C2A cancer cells following treatment with Ac$_5$GalNTGc, itraconazole, or P-3FAX-Neu5Ac. **d**, Cytotoxicity of unmodified NK-92 cells or HER2 CAR-expressing NK-92 cells against OE-19 cancer cells following treatment with Ac$_5$GalNTGc, itraconazole, or P-3FAX-Neu5Ac. Results are mean ± s.d. of n = 3 independent measurements. Statistical analysis in **a,c,d** was performed using one-way ANOVA with Tukey's multiple-comparisons test.

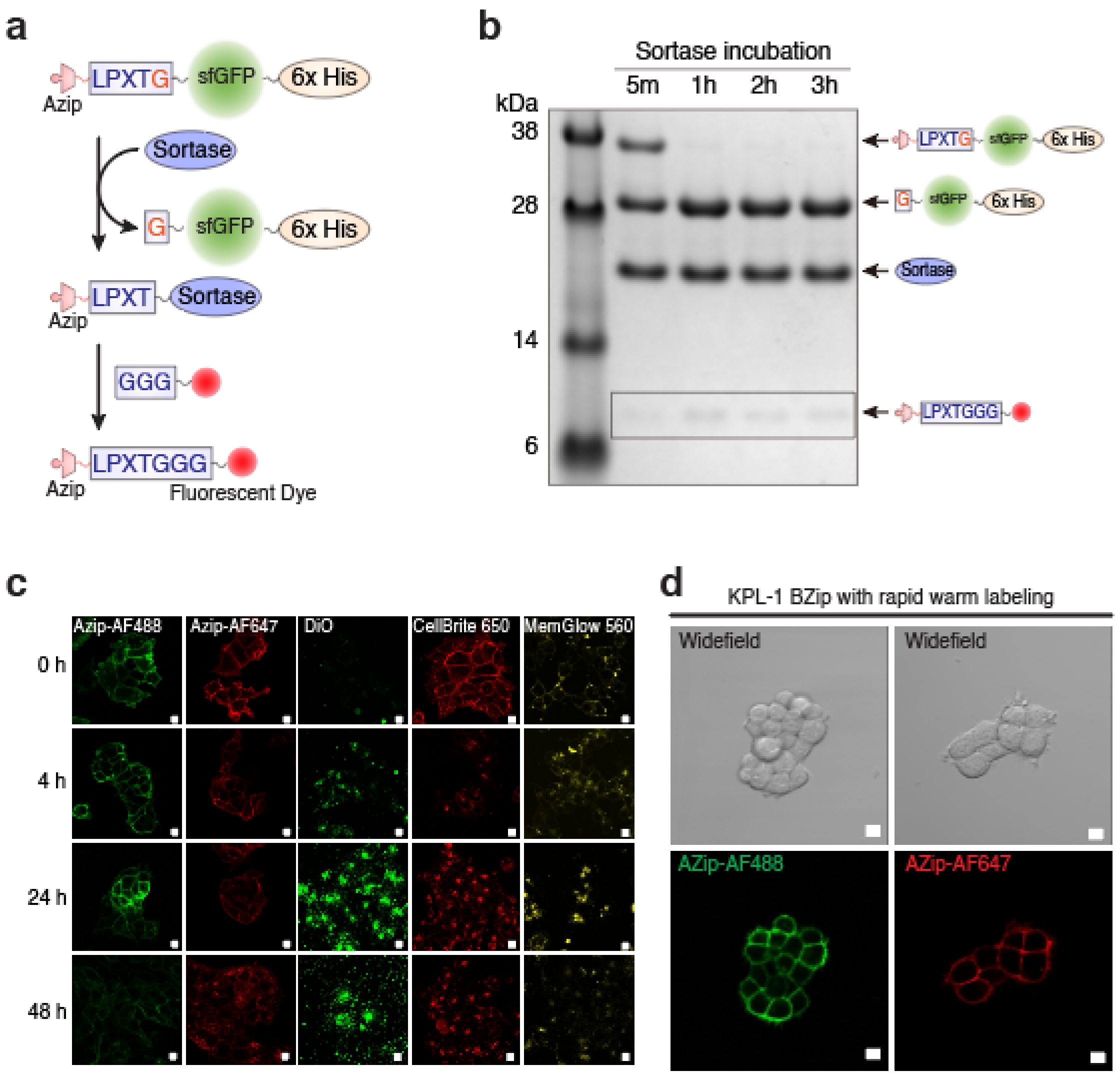


**Supplementary Figure 2. Sortase-mediated AZip fluorophore conjugation and membrane-label retention over time. a,** Schematic of sortase-mediated fluorophore conjugation to AZip-sfGFP. Sortase cleaves the LPXTG motif of AZip-sfGFP and catalyzes ligation to a GGG-functionalized fluorescent dye, generating fluorescently labeled AZip. **b**, Coomassie blue-stained gel showing the time course of the sortase-mediated reaction used to generate fluorescently labeled AZip. Reaction products corresponding to the indicated intermediates and final labeled products are shown. **c**, Representative fluorescence images of BZip-expressing KPL-1 cells labeled with AZip-AF488, AZip-AF647, DiO, CellBrite 650, or MemGlow 560 and imaged at 0, 4, 24, and 48 h after labeling. Cells were labeled for 10 min at 37 °C using the manufacturers' recommended concentrations and buffer conditions, followed by two washes with culture medium. These images show the complete time-course dataset corresponding to the membrane-labeling stability analyses presented in **Fig. 2g-i**. Scale bars, 10 µm. **d**, Leucine zipper-based membrane labelling with rapid warm condition**.** Fluorescence and bright-field images of Bzip-overexpressing KPL-1 cancer cells with 1 µM AZip-sfGFP for 10 minutes at 37°C instead of 1 hour at 4°C. Scale bars, 10 µm.

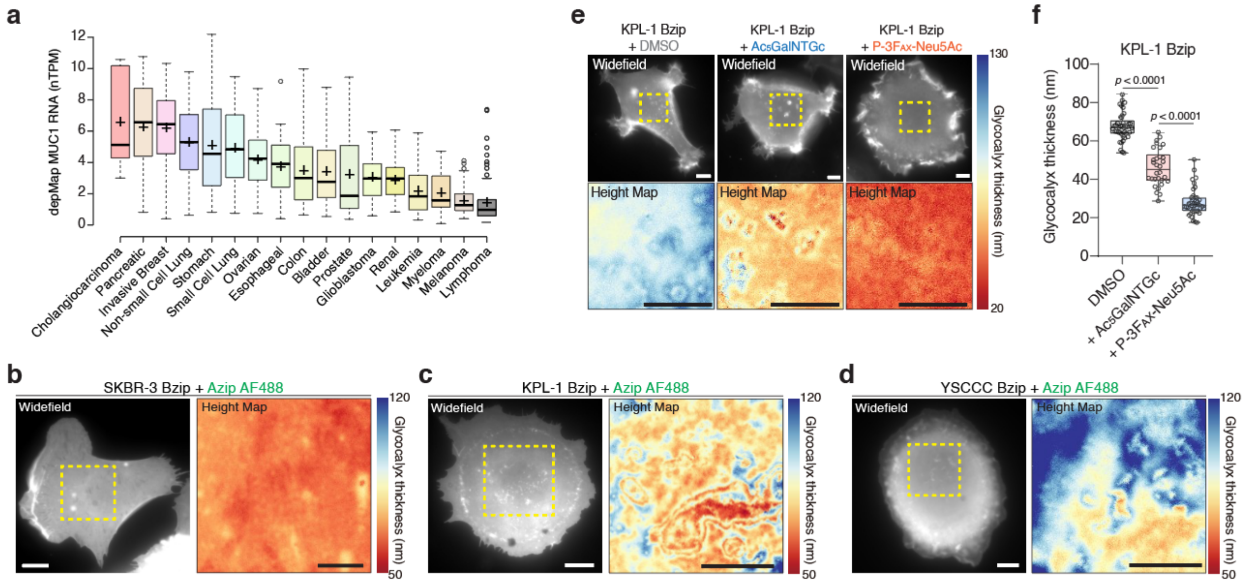


**Supplementary Figure 3. Validation of leucine zipper-based SAIM across cancer cell lines and pharmacological glycocalyx perturbations. a,** MUC1 transcript abundance across the indicated cancer types obtained from the DepMap dataset. Center lines indicate medians, "+" symbols indicate means, boxes indicate the 25th and 75th percentiles, whiskers extend to 1.5 times the interquartile range, and individual points outside the whiskers represent outliers. **b-d**, Representative wide-field fluorescence images and corresponding SAIM-derived glycocalyx height maps of live BZip-expressing SKBR3 (**b**), KPL-1 (**c**), and YSCCC (**d**) cells labeled with AZip-AF488 (1 μM) for 10 min at 37 °C. Scale bars, 10 μm. **e**, Representative wide-field fluorescence images and corresponding glycocalyx height maps of BZip-expressing KPL-1 cells following treatment with DMSO, 100 μM $Ac_5$GalNTGc, or 100 μM P-3$F_{AX}$-Neu5Ac for 48 h and subsequent labeling with AZip-AF488. Scale bars, 10 μm. **f**, Quantification of glycocalyx thickness in BZip-expressing KPL-1 cells following the indicated treatments. Boxes indicate the first and third quartiles, center lines indicate the median, and whiskers indicate the range. Each condition includes at least 30 cells from a representative experiment. Statistical significance was determined by one-way ANOVA followed by Tukey's multiple-comparisons test.

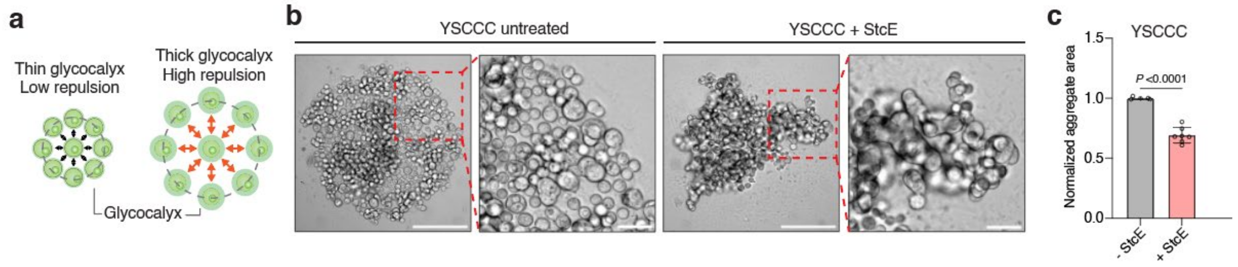


**Supplementary Figure 4. Glycocalyx reduction promotes multicellular aggregation in YSCCC cells.** **a,** Schematic illustrating the predicted effect of glycocalyx-dependent steric repulsion on multicellular aggregate formation. Cells with a thick glycocalyx experience greater cell-cell repulsion, whereas reduction of the glycocalyx is expected to facilitate closer cell-cell interactions and aggregate compaction. **b**, Representative bright-field images of multicellular aggregates formed by YSCCC cells cultured in round-bottom ultra-low-attachment wells in the presence or absence of 10 nM StcE for 24 h. A total of 1,000 cells were seeded per well. Enlarged regions corresponding to the dashed boxes are shown on the right. Scale bars, 200 µm; enlarged images, 50 µm. **c**, Quantification of aggregate area normalized to untreated YSCCC cells. Data are shown as mean ± s.d. for untreated (n = 5 aggregates) and StcE-treated (n = 7 aggregates) conditions. Statistical significance was determined using a two-tailed unpaired Student’s t-test.

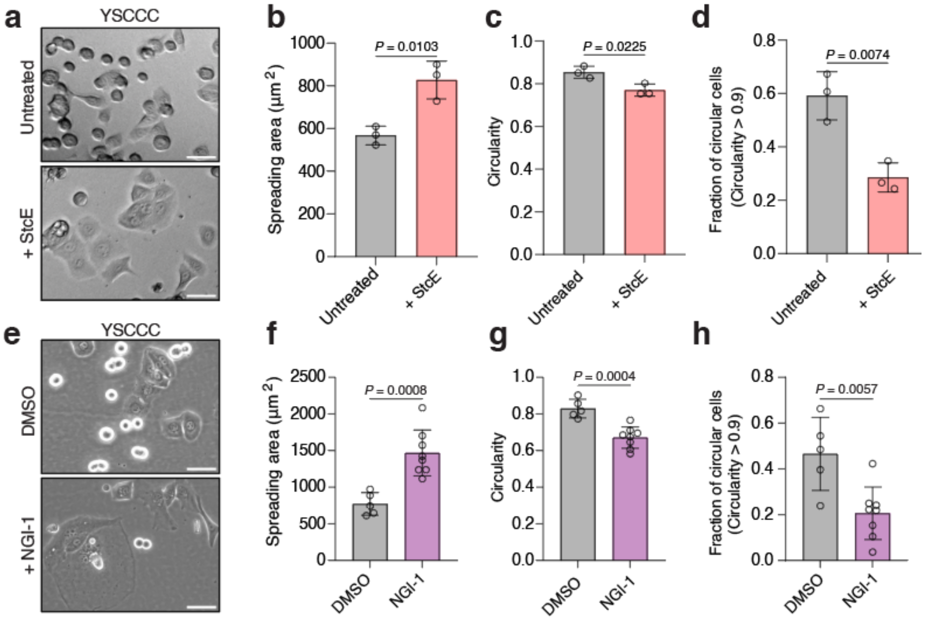


**Supplementary Figure 5. Glycocalyx perturbation alters cancer-cell spreading and morphology. a,** Representative phase-contrast images of YSCCC cells following overnight culture in the presence or absence of 1 nM StcE. Scale bars, 50 µm. **b-d**, Quantification of cell spreading area (**b**), circularity (**c**), and the fraction of highly circular cells (**d**) in untreated or StcE-treated YSCCC cells. Highly circular cells were defined as cells with a circularity value >0.9. Data are shown as mean ± s.d. from n = 3 fields of view, with approximately 100 cells analyzed per condition in each field. e, Representative phase-contrast images of YSCCC cells following overnight culture with DMSO or NGI-1. Scale bars, 50 µm. **f-h**, Quantification of cell spreading area (**f**), circularity (**g**), and the fraction of highly circular cells (**h**) in DMSO- or NGI-1-treated YSCCC cells. Highly circular cells were defined as cells with a circularity value >0.9. Data are shown as mean ± s.d. from n = 5-8 fields of view, with approximately 50 cells analyzed per condition in each field. Statistical significance in **b-d** and **f-h** was determined using two-tailed unpaired Student's t-tests.

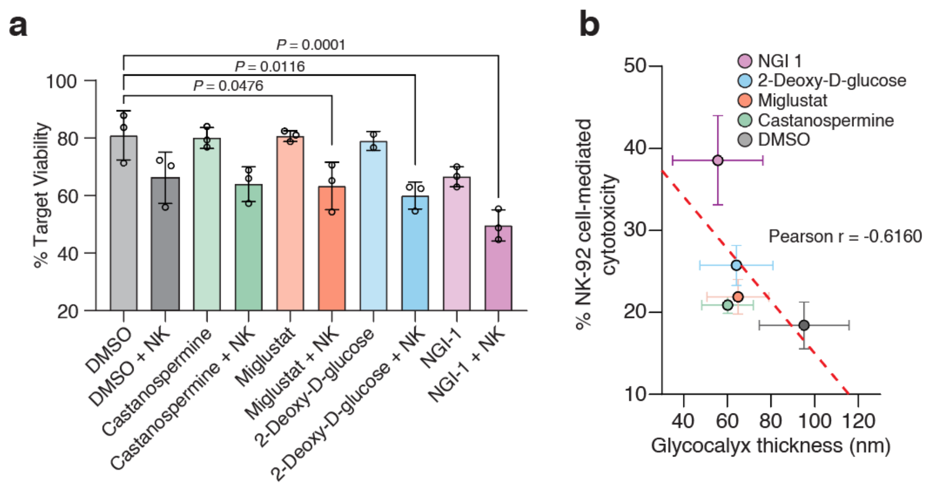


**Supplementary Figure 6. Relationship between pharmacological glycocalyx reduction and NK cell-mediated cytotoxicity. a,** Target-cell viability following treatment with the indicated pharmacological inhibitors in the presence or absence of NK-92 cells. Target cells were treated with castanospermine, miglustat, 2-deoxy-D-glucose, or NGI-1 and subsequently co-cultured with NK-92 cells. Target-cell viability was quantified relative to the corresponding untreated control. **b**, Relationship between glycocalyx thickness and NK-92 cell-mediated cytotoxicity following treatment with the indicated inhibitors. Glycocalyx thickness measurements are from **Fig. 4c**. Points represent mean ± s.d. for each treatment condition. The dashed line indicates the linear regression fit; Pearson r = -0.6160. In **a**, statistics were determined using a one-way ANOVA with Tukey's multiple comparisons test

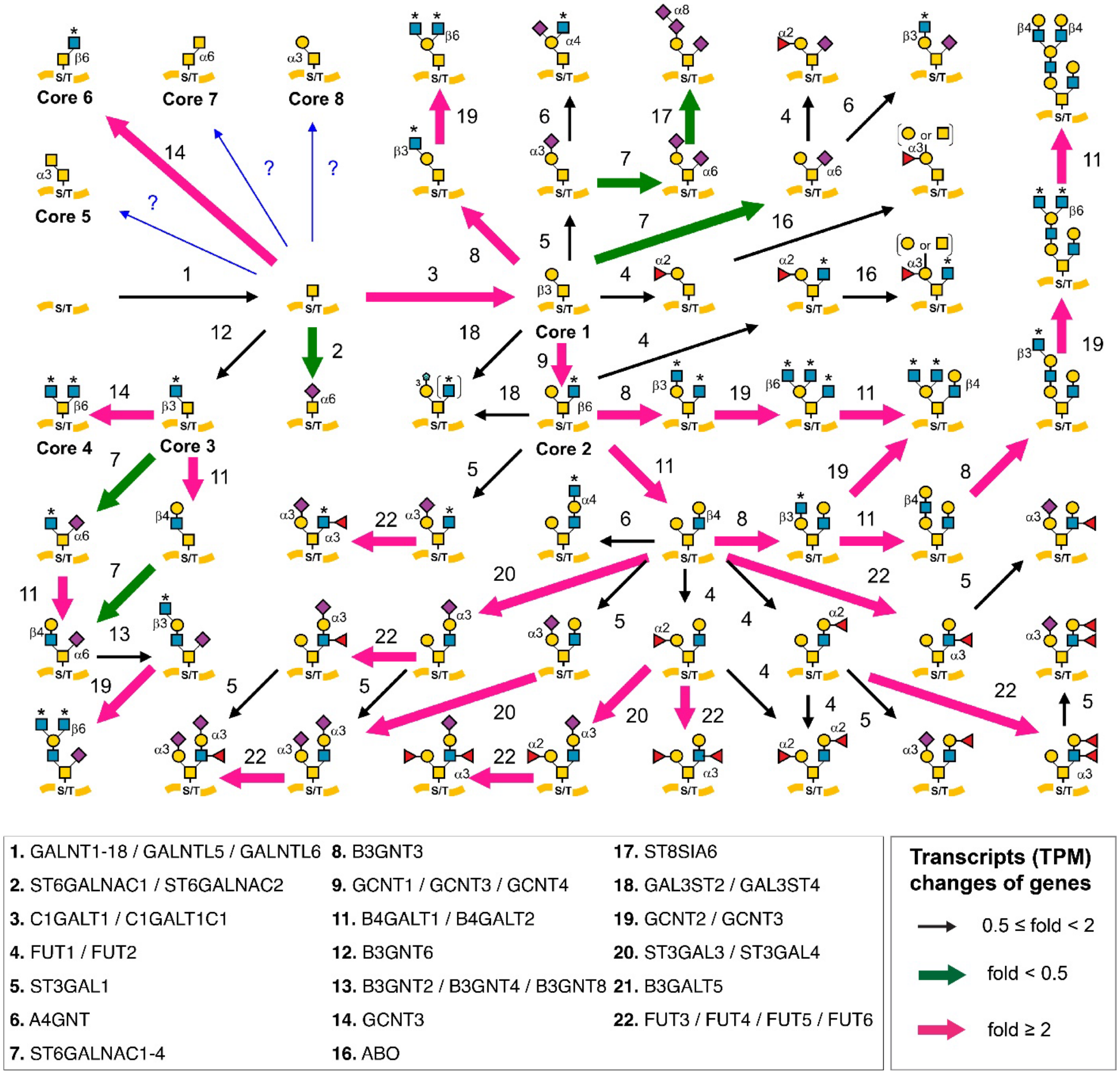


**Supplementary Figure 7. Comparative analysis of the biosynthetic pathways for mucin-type *O*-glycans between YSCCC and KPL-1 using GlycoMaple.** Reaction arrows are scaled and color-coded based on the normalized gene expression (transcripts per million; TPM) of the governing enzymes analyzed by RNA-Seq. If several enzymes have overlapping function in a reaction, the maximum TPM value among the values of overlapped genes was used. When several gene products make a complex for a reaction, the minimum TPM value of the subunit genes was used. Gene expression profiles (median TPM values) in YSCCC and KPL-1 were used to show the pathways. The fold change comparison is based on TPM values between YSCCC and KPL-1, where pink indicates cases where the fold change is greater than 2 (YSCCC is greater than KPL-1), and green indicates cases where it is less than 0.5 (KPL-1 is greater than YSCCC). Glycan structures are drawn using the standard Symbol Nomenclature for Glycans (SNFG). Transcriptomic data was from the Broad Cancer Cell Line Encyclopedia.

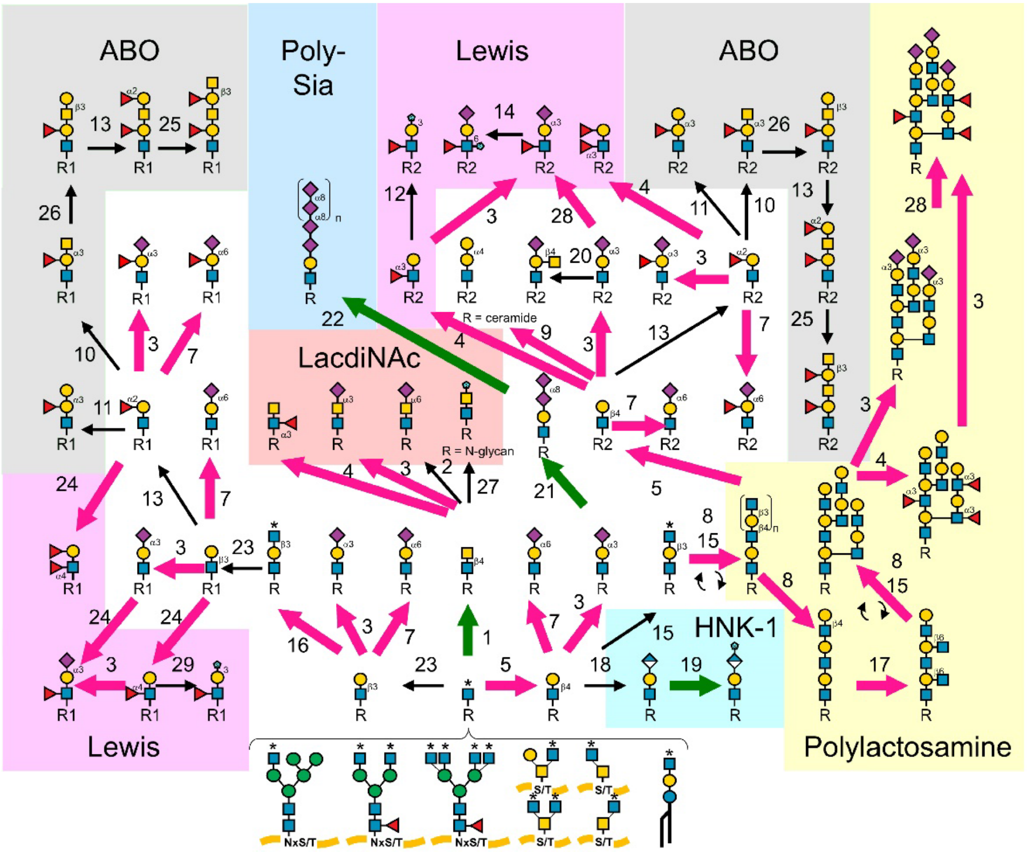


**1.** B4GALNT3 / B4GALNT4
**2.** ST6GAL2
**3.** ST3GAL3 / ST3GAL4 / ST3GAL6
**4.** FUT9 / FUT3 / FUT4 / FUT5 / FUT6
**5.** B4GALT1-5
**7.** ST6GAL1 / ST6GAL2
**8.** B4GALT1
**9.** A4GALT
**10.** ABO
**12.** GAL3ST2 / GAL3ST3
**13.** FUT1 / FUT2
**14.** CHST2 / CHST4
**15.** B3GNT2 / B3GNT4 / B3GNT8
**16.** B3GNT3
**17.** GCNT2 / GCNT3
**18.** B3GAT1 / B3GAT2
**19.** CHST10
**20.** B4GALNT2
**21.** ST8SIA2 / ST8SIA3 / ST8SIA4
**22.** ST8SIA2 / ST8SIA4
**23.** B3GALT1 / B3GALT2 / B3GALT5
**24.** FUT3
**25.** B3GALNT2
**26.** B3GALT1 / B3GALT2 / B3GALT5
**27.** CHST8 / CHST9
**28.** FUT7 / FUT3 / FUT4 / FUT5 / FUT6 / FUT9
**29.** GAL3ST2

**Transcripts (TPM) changes of genes**

→ 0.5 ≤ fold < 2
→ fold < 0.5
→ fold ≥ 2

**Supplementary Figure 8. Comparative analysis of the biosynthetic pathways for capping structures between YSCCC and KPL-1 using GlycoMaple.** Visualization of the biosynthetic pathway for capping structures in YSCCC and KPL-1 cells. The setting of arrows was described in **Extended Data Figure 7**. Gene expression profiles (median TPM values) in YSCCC and KPL-1 were used to show the pathways. The fold change comparison is based on TPM values between YSCCC and KPL-1, where pink indicates cases where the fold change is greater than 2 (YSCCC is greater than KPL-1), and green indicates cases where it is less than 0.5 (KPL-1 is greater than YSCCC). Glycan structures are drawn using the standard Symbol Nomenclature for Glycans (SNFG). Transcriptomic data was from the Broad Cancer Cell Line Encyclopedia.

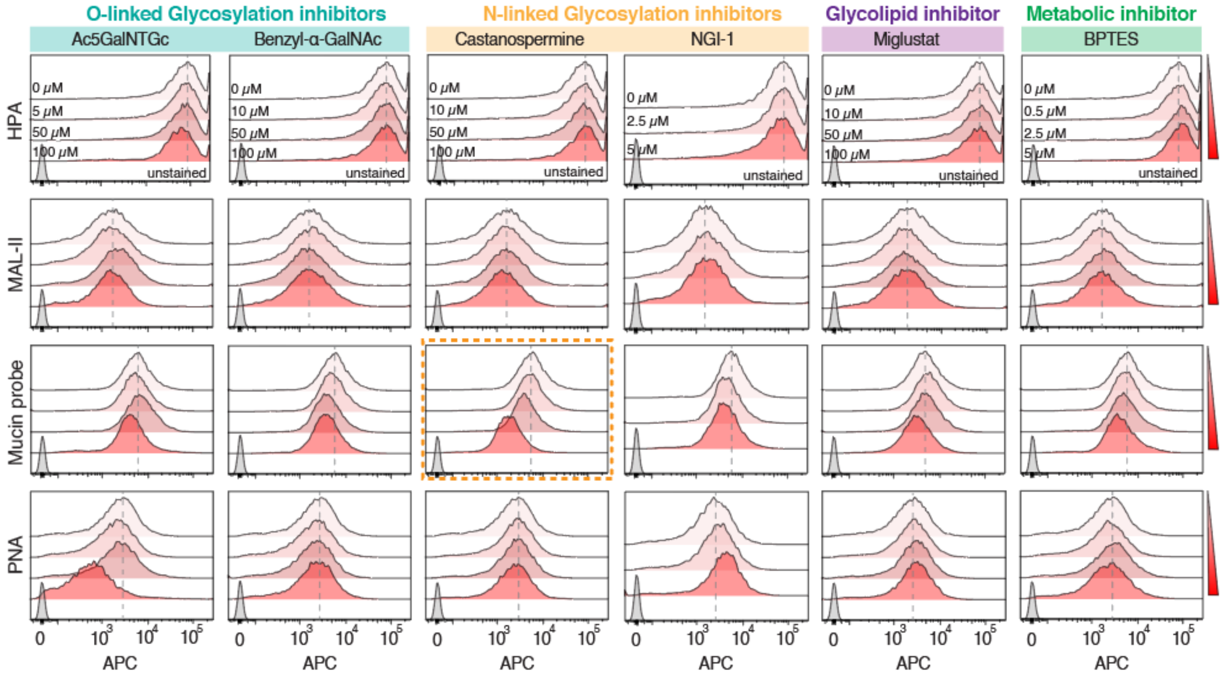


**Supplementary Figure 9. Flow cytometry analysis of glycocalyx remodeling by candidate pharmacological inhibitors in GSU cells.** Representative flow cytometry histograms showing HPA, MAL-II, mucin-probe, and PNA binding in GSU cells following treatment with the indicated concentrations of $Ac_5$GalNTGc, benzyl-α-GalNAc, Castanospermine, NGI-1, Miglustat, or BPTES. Untreated cells (0 μM) and unstained controls are shown for comparison. These measurements were used to quantify inhibitor-induced changes in cell-surface glycan and mucin features summarized in the heat map in **Fig. 4f**.

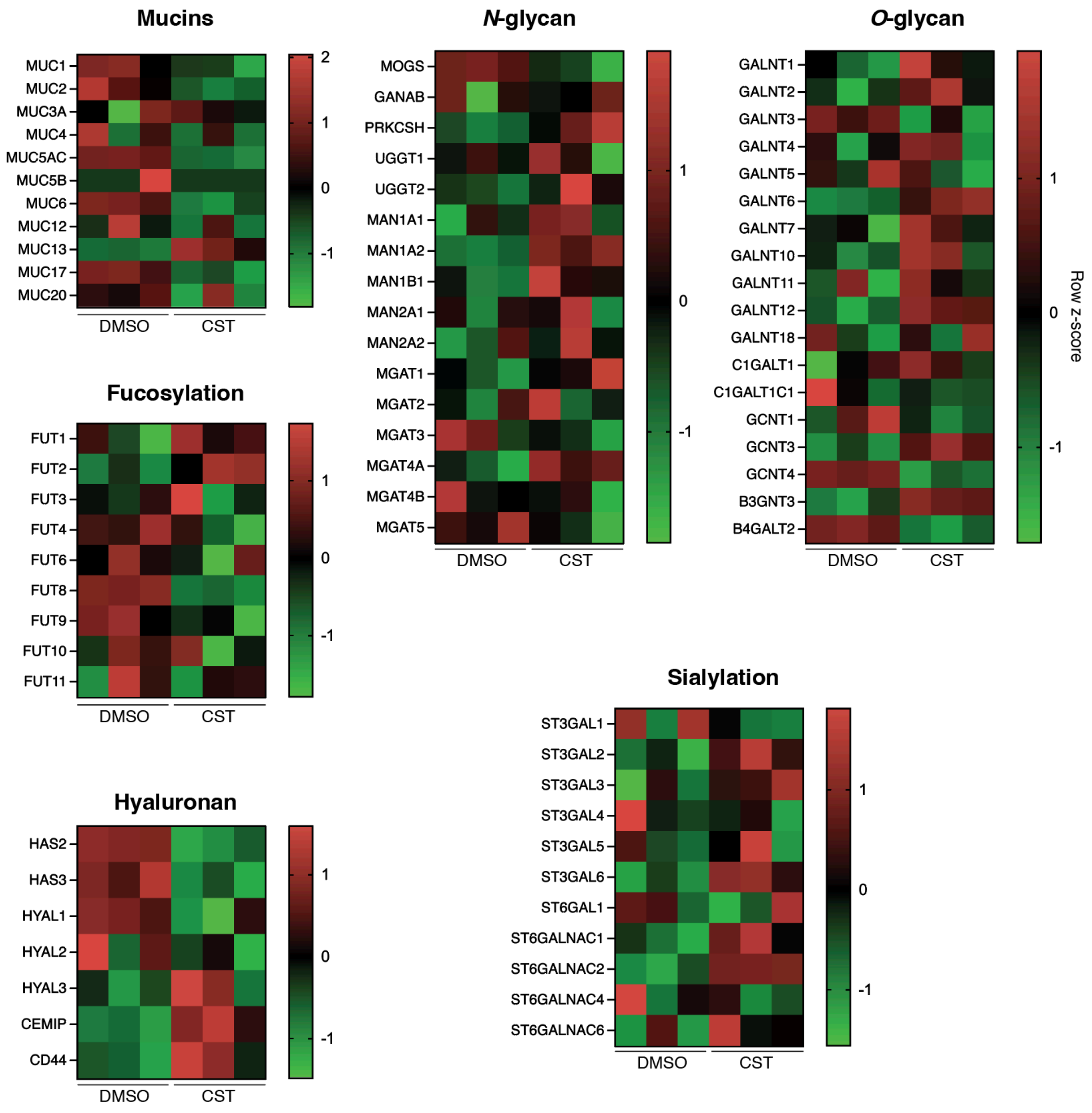


**Supplementary Figure 10.** Heatmap of glycocalyx-associated genes in DMSO- and Castanospermine (CST)-treated GSU cells, grouped according to mucin, hyaluronan, *N*-glycan, *O*-glycan, sialylation, fucosylation-associated functions from **Fig. 5c**. Each row represents an independent biological replicate. Gene expression values were normalized as counts per million (CPM), $\log_2$-transformed, and displayed as row-wise z-scores.

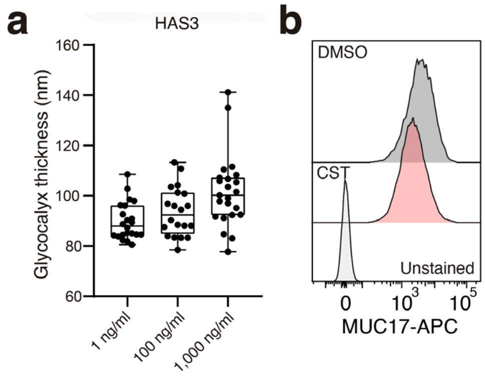


**Supplementary Figure 11.** ***HAS3*** **expression contributes to glycocalyx thickness, whereas castanospermine does not increase cell-surface MUC17 expression.** **a**, Quantification of glycocalyx thickness in cells carrying a doxycycline-inducible *HAS3* expression system following treatment with 1, 100 or 1,000 ng/ml doxycycline. Boxes indicate the first and third quartiles, center lines indicate the median, and whiskers indicate the range. **b**, Representative flow cytometry histograms showing cell-surface MUC17 staining in GSU cells treated with DMSO or castanospermine (CST). Unstained cells are shown as a negative control.

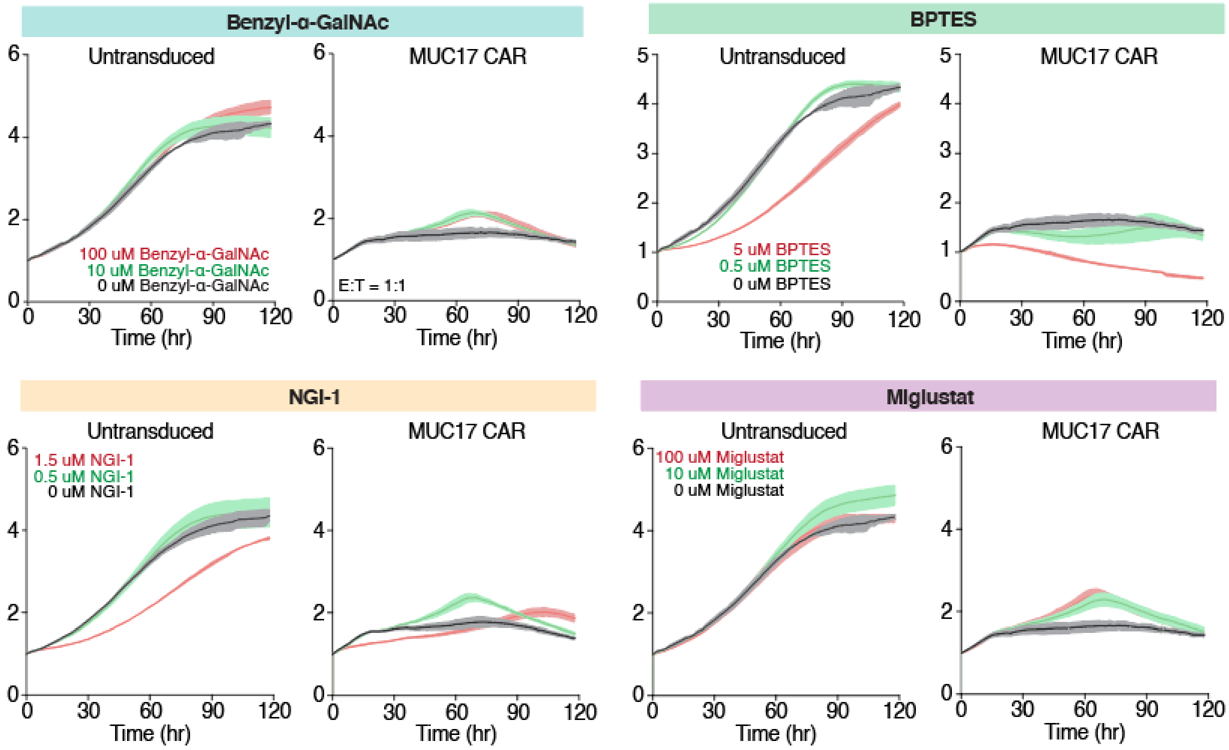


**Supplementary Figure 12. Functional evaluation of additional glycocalyx-modulating compounds in MUC17 CAR-T cell cytotoxicity assays.** Real-time cytotoxicity assays of GSU target cells co-cultured with untransduced T cells or MUC17 CAR-T cells following treatment with the indicated concentrations of benzyl-α-GalNAc, BPTES, NGI-1, or miglustat. Cancer cell growth was monitored by normalized GSU GFP fluorescence intensity over time. MUC17 CAR-T cells were co-cultured with GSU target cells at an E:T ratio of 1:1. The corresponding castanospermine experiment is shown in **Fig. 5f**.
